# Systems immunology of transcriptional responses to viral infection identifies conserved antiviral pathways across macaques and humans

**DOI:** 10.1101/2023.06.22.546003

**Authors:** Kalani Ratnasiri, Hong Zheng, Jiaying Toh, Zhiyuan Yao, Veronica Duran, Michele Donato, Mario Roederer, Megha Kamath, John-Paul M. Todd, Matthew Gagne, Kathryn E. Foulds, Joseph R. Francica, Kizzmekia S. Corbett, Daniel C. Douek, Robert A. Seder, Shirit Einav, Catherine A. Blish, Purvesh Khatri

## Abstract

Viral pandemics and epidemics pose a significant global threat, with emerging and re-emerging viruses responsible for four pandemics in the 21st century alone. While macaques have been utilized as a model for understanding viral disease in a controlled setting, it remains unclear how conserved the antiviral responses to diverse viruses are between macaques and humans. To address this critical knowledge gap, we conducted a comprehensive cross-species analysis of transcriptomic data from over 6000 blood samples from macaques and humans infected with one of 31 viruses, including Lassa, Ebola, Marburg, Zika, and dengue. Our findings demonstrate that irrespective of primate or viral species, there are conserved antiviral responses which are consistent regardless of infection phase (acute, chronic, or latent) and viral genome type (DNA or RNA viruses). Moreover, by leveraging longitudinal data from experimental challenges, we identified virus-specific response dynamics such as host responses to Coronaviridae and Orthomyxoviridae infections peaking 1-3 days earlier than responses to Filoviridae and Arenaviridae viral infections. Additionally, through comparative analysis of immune responses across viruses, we identified a unique enrichment of lymphoid cellular response modules in macaque Flaviviridae infection that persists in human responses to dengue. Our results underscore macaque studies as a powerful tool for gaining new insights into viral pathogenesis and immune responses that translate to humans, which can inform viral therapeutic development and enable pandemic preparedness.

**One sentence summary:** Using longitudinal macaque viral challenge studies, we identified shared and virus-specific responses to infection that replicate in human viral disease - thereby demonstrating the utility of macaque models of viral infection to understand antiviral biology and for pandemic preparedness.

## Introduction

Current, emerging, and reemerging viruses constantly threaten human health, not only by causing disease and death, but also by driving wider societal and global consequences. Estimates suggest that RNA viruses make up to 44% of all emerging infectious diseases, with 2-3 novel virulent viruses discovered yearly and most of zoonotic origins.^1, 2^ Particular RNA viral families, including Flaviviridae, Coronaviridae and Orthomyxoviridae, have led to multiple epidemics and pandemics within the 21st century^3^, demonstrating their pandemic potential. RNA viruses constantly evolve: mistake-prone RNA polymerases introduce genomic mutations and zoonotic reservoirs drive unique evolutionary pressures on viruses that lead to unpredictable emergence patterns and disease manifestations.^4, 5^ While controlled human infection studies are ideal for developing translational solutions, such studies are generally difficult and unethical for lethal and emerging pathogens. Therefore, non-human primate (NHP) models, particularly the macaque model, continue to be critical for understanding disease pathogenesis, vaccine modalities, and therapeutic interventions.^6^

Previously, we identified a conserved host response in human infection across multiple viruses that have led to epidemics and pandemics.^7, 8^ However, multiple questions remain. For example, determining generalizability of host responses across infection by viruses such as Marburg and Lassa is needed; yet, the lack of available data on human infections caused by these and other viruses impedes the assessment of panviral responses. Additionally, understanding early antiviral responses remains important though complicated in human profiling studies due to the challenge of identifying time-of-infection and virus incubation periods. Furthermore, while it is necessary to compare the longitudinal dynamics of host response induction across viruses, ethical concerns exist regarding human viral challenge studies. Here, macaque studies are advantageous because they allow for the understanding of diverse and lethal viral pathogens in well-controlled challenge studies, whereby measurements can be taken across multiple timepoints pre- and post-infection. However, the extent to which macaque immune responses reflect human host responses or vice versa is unclear, particularly whether both humans and macaques evoke similar antiviral responses upon infection. By leveraging transcriptomic profiles from both macaque and human infection studies, we aim to determine the utility of macaque models for understanding and predicting human responses to emerging viruses and to map conserved and unique features of the immune response to different viruses.

In this study, we performed the largest transcriptome analysis of viral disease in macaques and humans to (1) directly compare human and macaque antiviral responses and (2) define conserved and unique features of host responses across viruses of concern. We used blood transcriptome data from 21 bulk RNA-seq datasets comprising 743 samples from 198 macaques from three species of macaques (rhesus, cynomolgus and pigtailed macaques) and infection by 13 viruses across five viral families. We utilized longitudinal data to analyze the dynamics of viral response induction across numerous viruses, some of which have been seldom studied in the context of human transcriptomic responses. Further, we applied our previously identified conserved human host response, Meta-Virus Signature (MVS), which distinguishes viral infection from healthy controls and predicts severity in humans, to show that macaques also induce antiviral responses similar to those of humans and that these response dynamics vary by viral family.^7, 8^ We also demonstrate that conserved responses in NHP data robustly translate to human transcriptomic responses to heterogenous viral infections by leveraging 5345 human samples across 47 datasets. Additionally, comparative analysis across antiviral responses allows us to identify differentiating features of T cell responses in Flaviviridae infection of macaques that replicate in human studies. Together, this work demonstrates that macaque transcriptomic antiviral responses robustly recapitulate those in human viral disease and are conserved across diverse viruses - further supporting the use of macaque models to develop antiviral countermeasures, particularly in cases where human studies are not possible.

## Results

### Data collection, curation, and preprocessing

We searched public repositories and publications for blood transcriptomic datasets from macaques with viral infection. We focused on acute RNA viruses from the World Health Organization (WHO) list of priority pathogens.^9^ We also included Orthomyxoviridae due to its history of, and potential for, driving pandemics (Tables 1 and S1). We identified 21 datasets composed of 743 samples from 198 macaques infected with one of 13 viruses across five viral families (Tables 1 and S1). Together, these datasets represented a broad spectrum of biological and technical heterogeneity as they included data from three different macaque species infected with one of 13 viruses via different routes and doses, and profiled using different microarray platforms and by RNA sequencing. Because timepoints across datasets were not uniform, we grouped timepoints into six discrete categories, where T0 included uninfected samples prior to challenge, T1 spanned days 1-2 post-infection, T2 was days 3-5, T3 was days 6-8, T4 was days 9-13 and T5 was days 14+ (Figure 1A). While most datasets included animals from baseline through infection, one Arenaviridae challenge dataset did not have pre-infection timepoints, requiring unpaired analyses when this dataset was included. Before analyzing all the macaque species together, we confirmed that the macaque species were comparable at baseline by comparing pairwise correlation of the 2,083 shared genes across the different datasets. We found that there were no differences across datasets of different macaque species when compared to the variation seen across datasets within the same macaque species (Figure S1).

**Figure 1.**
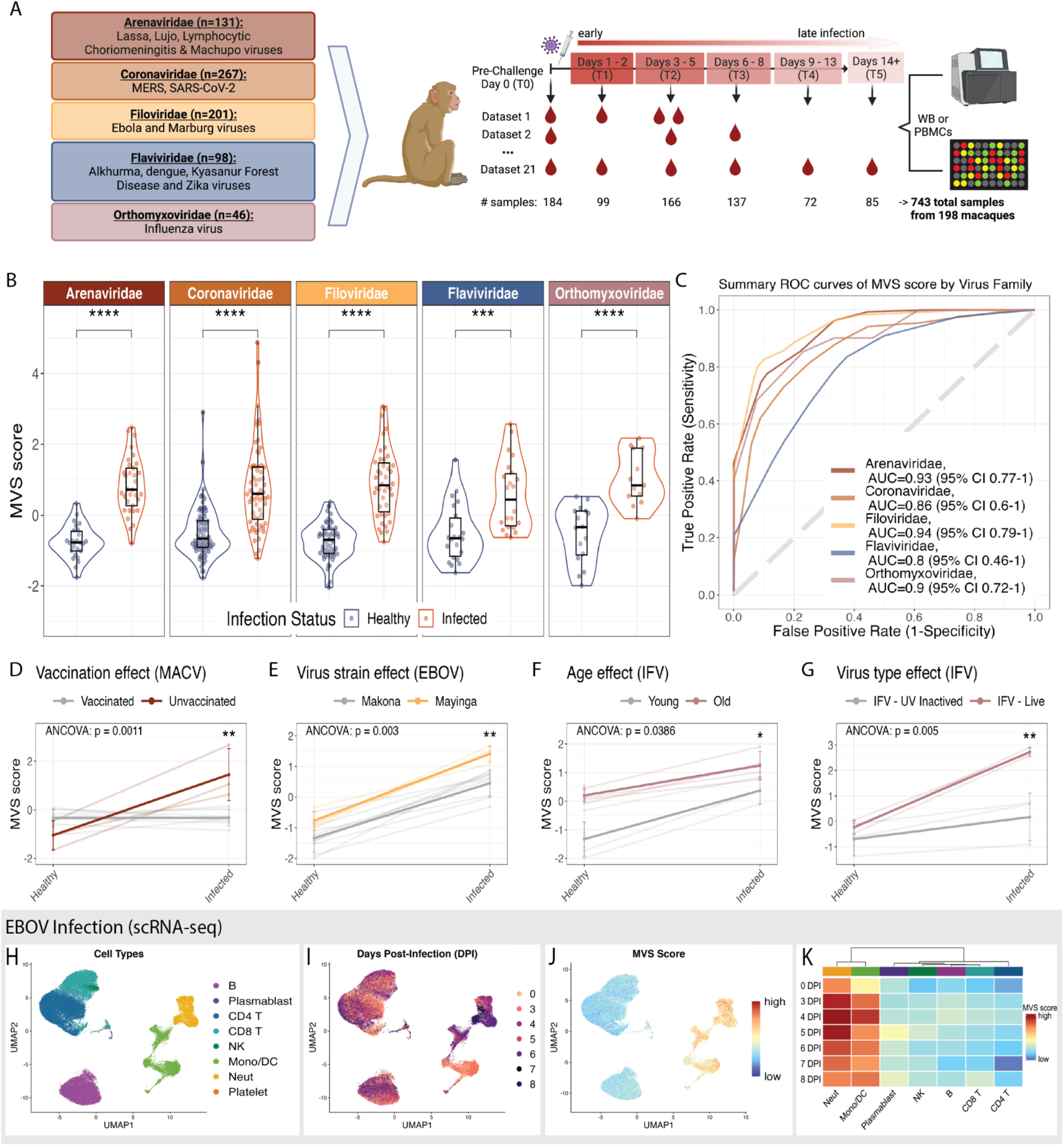
Conserved immune response to viral infection in humans is similarly present in macaque infection and driven by myeloid cells **(A)** Schematic of macaque sample overview and timepoint distribution. **(B)** Distribution of the Meta-Virus signature (MVS) scores comparing uninfected, healthy macaques to those at peak MVS score by viruses across five viral families. Each point represents a blood sample. Significance values were determined using an unpaired Wilcoxon ranked-sum test with Bonferroni correction for multiple hypothesis testing. **(C)** ROC curves for distinguishing macaques with viral infection at peak MVS timepoint category from uninfected macaques, colored by the viral family associated with infection (382 samples in 21 datasets). **(D-G)** Association of MVS scores with the known risk factors of disease severity **(D)** vaccination status, **(E)** virus strain, **(F)** age of host, and **(G)** live virus across four different datasets from macaques infected with Machupo, influenza or Ebola virus. P-value was determined by ANCOVA test accounting for MVS score at pre-infection timepoint and a risk factor of interest as a covariate of the MVS score post-infection. **(G-I)** UMAP visualization of 56,929 immune cells from 17 animals colored by **(H)** cell type, **(I)** day post-infection, and **(J)** MVS score. **(K)** Heatmap representing the average MVS score of each cell type across pre-infection and each day post-infection. Asterisk values across figure are represented as follows: *p-value < 0.05, **p-value < 0.01, ***p-value < 0.001, and ****p-value < 0.0001. WB = Whole blood, PBMC = Peripheral blood mononuclear cells, MACV = Machupo virus, EBOV = Ebola virus, IFV = influenza virus.

**Table 1:**
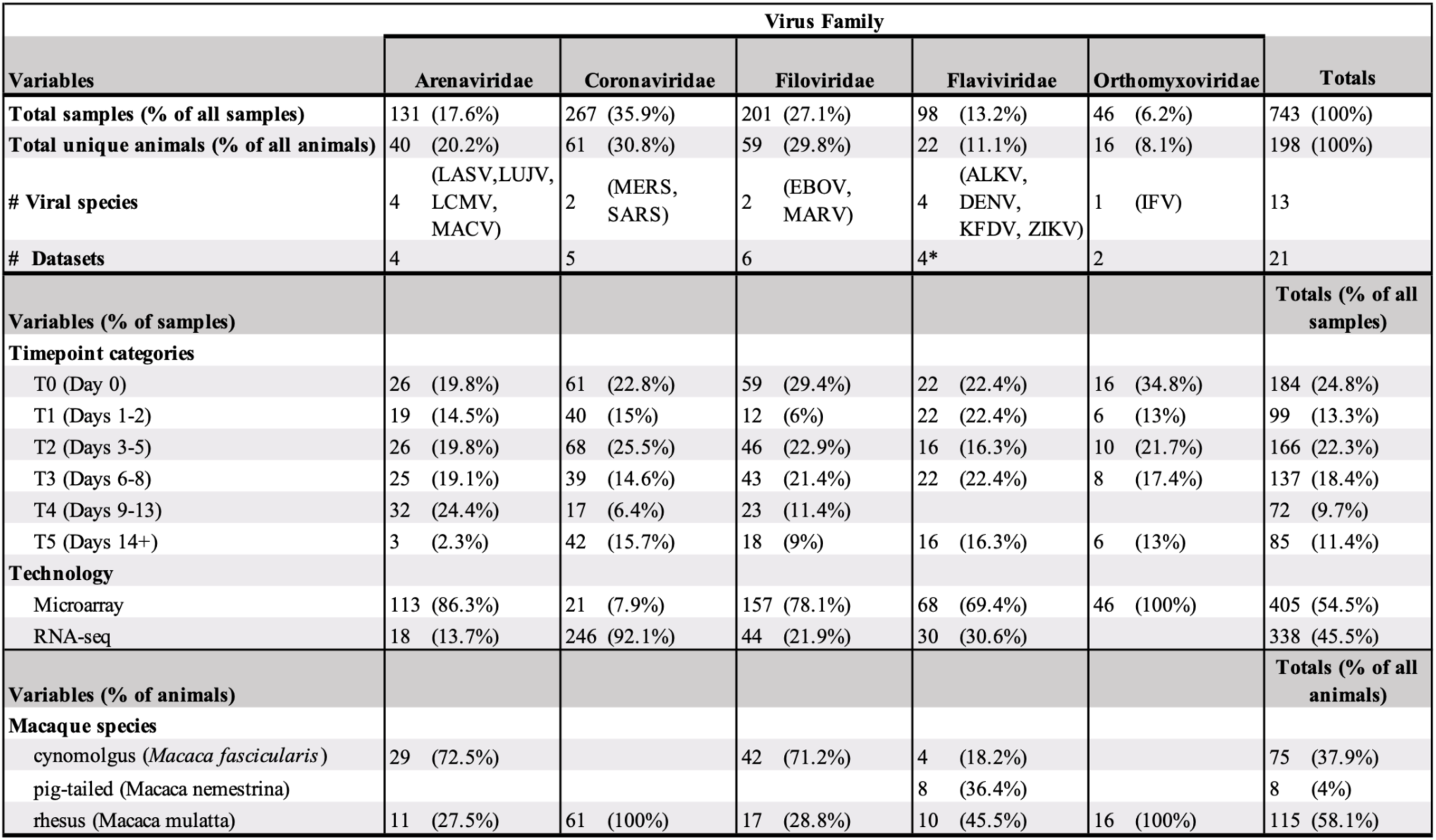
Sample distribution of macaque bulk RNA-seq datasets.

### Conserved immune response to viral infections in humans is also conserved in macaques to diverse RNA viruses

We asked whether macaques are a representative model for studying human immune responses to viral infections. To answer this question, we used the MVS, the conserved immune response signature we have described and validated previously.^7, 8^ As described before,^7^ we calculated the MVS score for each macaque sample in each dataset and compared the MVS scores at peak infection timepoint to those at the baseline. We defined peak infection timepoint in a dataset as the timepoint category with the highest median MVS score. The MVS scores were significantly higher (padj<0.001) and accurately distinguished macaques at peak infection from uninfected timepoints (area under the receiver operating characteristic (AUROC) curve >= 0.8) across all viruses (Figures 1B, 1C and S2). We also examined other gene sets previously demonstrated to correlate with human viral infection. All viral infection datasets showed significant increase in interferon- stimulated gene (ISG) expression (padj<0.001), whereas Arenaviridae, Coronaviridae, and Filoviridae infections demonstrated significant downregulation of HLA Class II genes (padj<0.05) and significant upregulation of MS1 signature genes (padj<0.01)^10^(Figures S3A, S3B and S3C).

Importantly, we have shown that the MVS score is significantly correlated with severity of viral infection in humans.^8^ There were four datasets with known risk factors for severity in macaques. The MVS score was significantly associated with risk factors (p<0.04) including: Machupo virus infection of unvaccinated (more severe disease) versus vaccinated macaques^11^ (Figure 1D), infection by Mayinga (more severe disease) versus Makona Ebola strains^12, 13^ (Figure 1E), influenza infection of old (more severe disease) versus young macaques^14^ (Figure 1F), and infection by live (more severe disease) versus inactivated influenza infection^15^ (Figure 1G). These results demonstrate that conserved transcriptional signatures to viral infections in humans are also conserved in macaques and associated with known risk factors for severe outcome.

Furthermore, we have previously found that the MVS is driven by myeloid cells in humans with COVID-19.^8^ Therefore, we analyzed a single-cell RNA-seq (scRNA-seq) dataset of whole blood samples from macaques infected with Ebola virus (56,929 cells from 17 macaques)^16^ to investigate whether the conserved immune response in macaques is driven by the same immune cells (Figures 1H and 1I). We found that, similar to humans with COVID-19, the MVS genes were preferentially expressed in myeloid cells^8^ (Figures 1J and 1K). These results further suggest that myeloid cells are also a major driver of the MVS for multiple viruses in macaques. To further characterize infection-driven changes in myeloid cells, we examined longitudinal gene expression profiles from pre-infection to 8 days post-infection when macques developed severe and fatal disease. We observed an increase in ISG expression (r=0.93, p<1e-4; Figure S3D), MS1 signature genes (r=0.71, p<1e-4; Figure S3E), and downregulation of HLA Class II genes (r=-0.61, p<6e-4; Figure S3F); these changes were also observed in myeloid cells in the setting of severe COVID-19.^10^

Additionally, upregulation of MS1 genes and downregulation of HLA Class II is consistent with the acquisition of a myeloid derived suppressor cell (MDSC)-like phenotype, which in turn suppresses T cell activation. Therefore, we evaluated changes in T cell activation across infection. We observed significant downregulation of T cell activation genes in both CD4 and CD8 T cell subsets (r <-0.5, p<0.007; Figures S3G and S3F) from pre-infection to day 8 post-infection. Together, these results reveal that the dynamics described in human COVID-19 are also present in critical/fatal Ebola infection of rhesus macaques, further supporting our hypothesis that immune cell responses are consistent across viral and host species in RNA virus infections.

Together, these data provide strong evidence that the conserved immune response to viral infections in humans is also conserved in macaques and primarily driven by myeloid cells, and further suggests that it may be correlated with severity of infection in macaques.

### Temporal patterns of the conserved antiviral responses differ by viral families in humans and macaques

Because the peak infection timepoint differed for each virus, we investigated whether temporal patterns of the host response differed by virus in humans and macaques. First, we identified seven human challenge studies (GSE73072) where participants were inoculated with either influenza (IFV; family: Orthomyxoviridae), human rhinovirus (HRV; family: Picornaviridae), or respiratory syncytial virus (RSV; family: Pneumoviridae) and transcriptional data was collected from blood samples across pre- and post-infection. We excluded participants that were asymptomatic and did not shed virus (i.e., uninfected). We calculated the MVS score at all timepoints collected in symptomatic infected patients (Figures 2A and S4A) and assessed temporal changes in the MVS score with different viral infections (Figure 2B; Table S3). While IFV and HRV infections had highest MVS scores between days 1-5 post-infection, RSV infection showed peak MVS scores at later timepoints, between days 3-7 (Figures 2A, 2B and S5). A mixed-effects model with time as a continuous variable also showed that dynamics of the MVS in RSV-infected patients differed significantly (p<0.001) from those of patients with IFV or HRV infections (Table S3).

**Figure 2:**
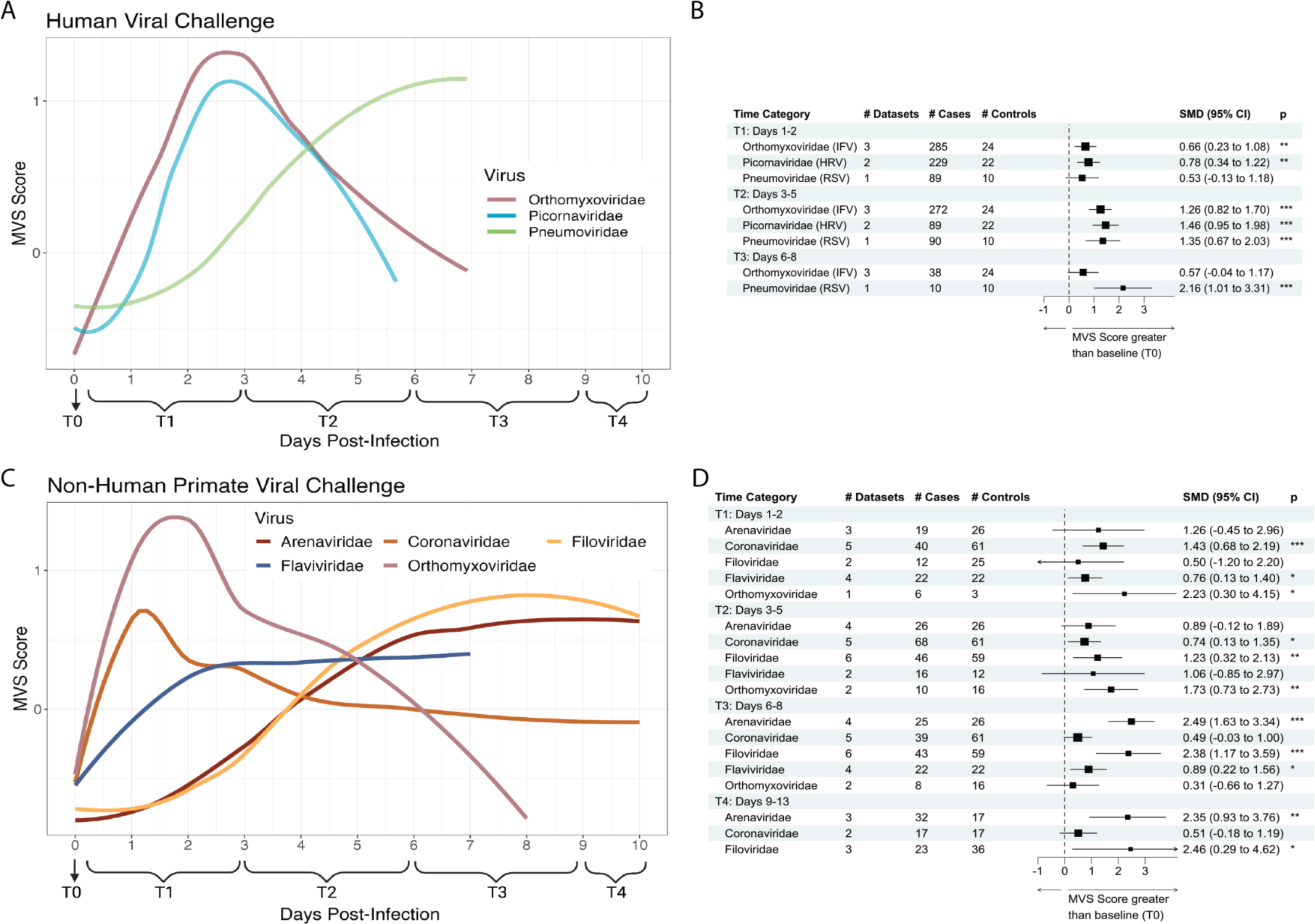
Longitudinal dynamics of the conserved antiviral response differ between viruses. MVS scores across all datasets up to 10 days post-infection across **(A)** 1158 human and **(C)** 734 NHP challenge samples with time category annotated below. Forrest plot tables of the summary statistics generated for each viral infection in **(B)** human and **(D)** NHP challenge dataset by time category.

Next, we investigated whether similar virus-dependent differences in the dynamics of the MVS were also present in macaques (Figures 2C and S6). Similar to IFV infection in humans, Orthomyxoviridae infection of macaques had early peak MVS responses at 1-3 days post infection (Figures 2C and 2D). We further comparatively assessed response dynamics via mixed-effects modeling using macaque infection by Orthomyxoviridae viruses as the comparator; however, we limited our analysis to pre-infection to day 7 post infection as Flaviviridae datasets had no timepoints past day 7 (Table 2). Dynamics of the MVS response in macaque infection by Arenaviridae and Filoviridae viruses were significantly different from those by Orthomyxoviridae infection (p<0.01), whereas infection by Coronaviridae and Flaviviridae viruses was less significant but did differ from Orthomyxoviridae infection dynamics (p<0.05; Table 2). For example, while Orthomyxoviridae and Coronaviridae only showed significant differences in MVS Scores at T1 and T2 compared to baseline (p<0.05), Filoviridae and Arenaviridae infections showed the most significant differences in MVS score compared to baseline at T3 (p<0.001; Figure 2D). Interestingly, both Filoviridae and RSV fall into the viral order Mononegavirales, and here we see both virus infections driving delayed peak MVS response induction in humans and macaques respectively (Figure 2). Overall, these data highlight that MVS is robustly conserved in humans and macaques across viruses, though host response dynamics differ by virus type that may be important for understanding viral incubation and latency periods.

**Table 2 :**
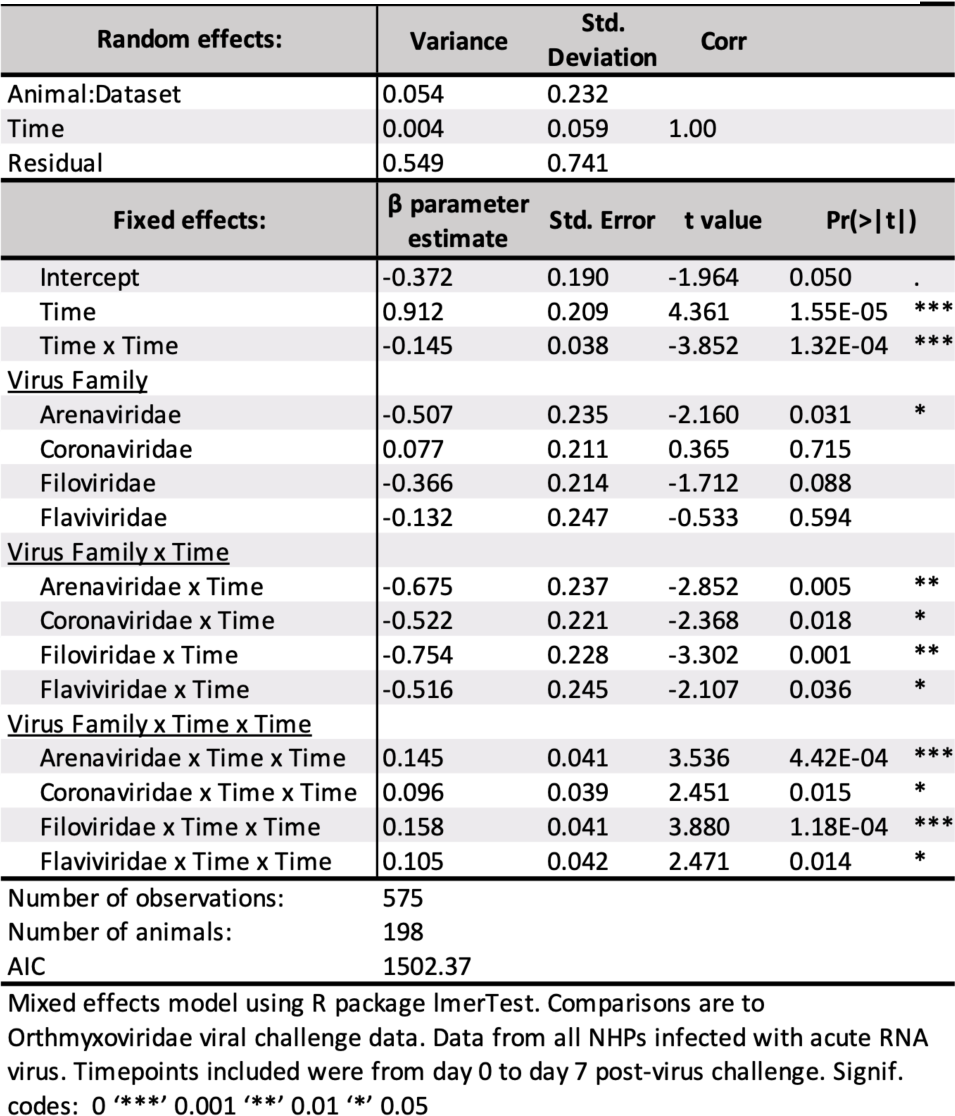
Time Series Analysis of MVS Score by Viral Challenge in NHPs

### Unbiased transcriptomic analysis of NHP demonstrates conserved antiviral responses to acute RNA viruses that translate to humans

Transcriptomic data for certain viral diseases (e.g., Lassa virus, Machupo virus, Kyasanur Forest Disease virus) in humans are not publicly available and other measurements at early stages of lethal and newly emerging viruses are likely lacking. In these cases, macaque studies provide the only immediate transcriptomic data to learn primate immune responses. Therefore, we investigated whether macaque antiviral responses were similarly conserved across viral infection and translatable to human viral infections (Figure 3A). First, we determined differentially expressed genes (DEGs) across datasets per viral family by timepoint category compared to the baseline T0 to identify peak infection timepoints in an unbiased way (Figure 3B). From here, we categorized peak infection timepoints per viral family as the time category with the highest number of robustly changing genes, defined as genes with false discovery rate (FDR)<0.05 and abs(effect size (ES))>0.1. Generally, peak DEG timepoint categories for each viral family corresponded with the peak MVS score timepoints demonstrated in Figure 1 (Figure 3B and Table S1).

**Figure 3:**
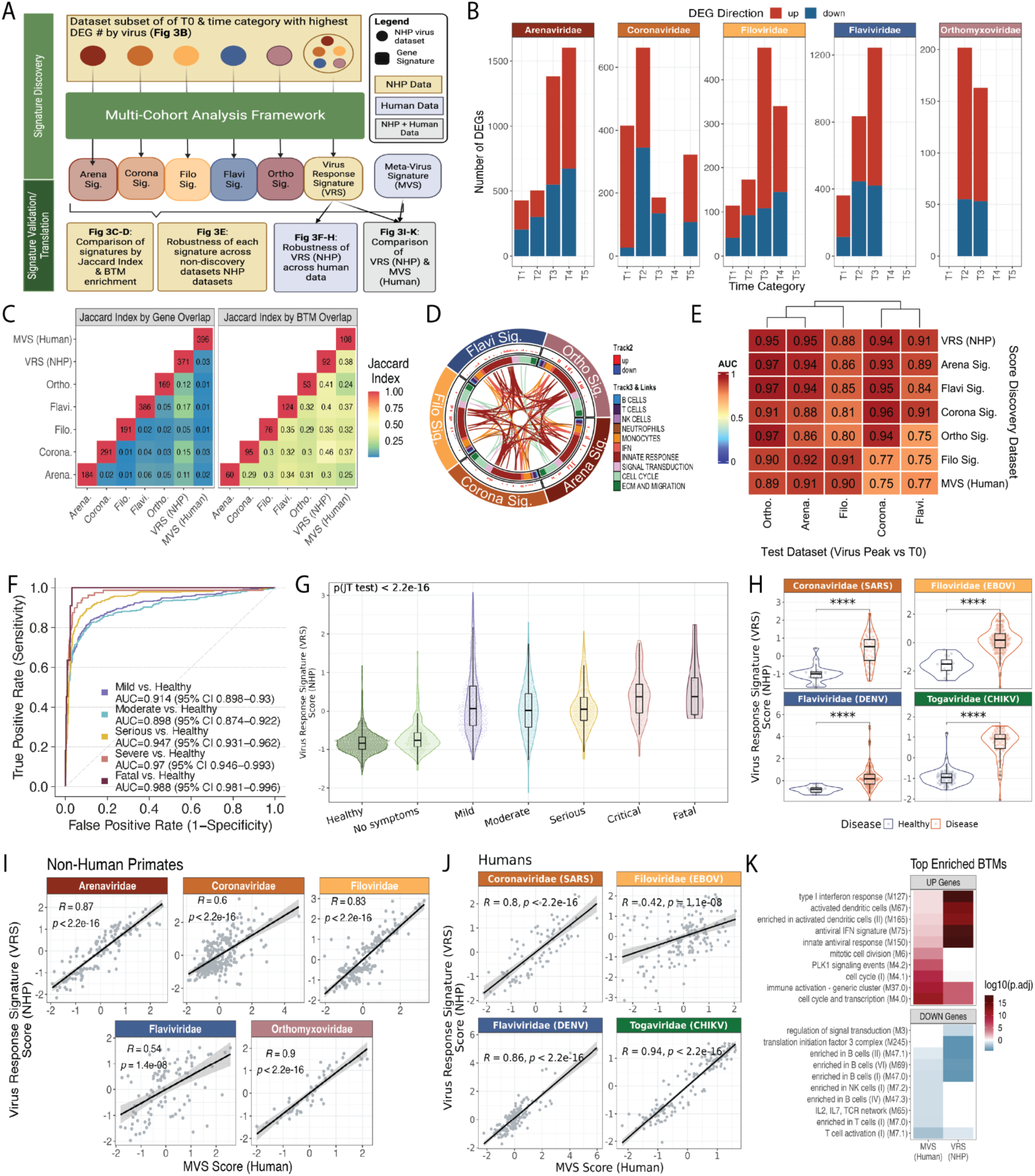
NHPs demonstrate conserved responses to acute RNA viruses that robustly translate to humans. **(A)** Schematic of experimental design for (B - K). **(B)** Significant DEGs at each timepoint category by viral family (effect size (ES) FDR < 0.05 and abs(ES) >0 .1). **(C)** Jaccard similarity index of the signature genes between each signature where annotation across the diagonal denoting same-score comparison is annotated with the number of genes present in the signature and all other annotations are the calculated Jaccard index (left). Jaccard similarity index of the blood transcription modules (BTM) that contain one or more of the signature genes between each signature where annotation across the diagonal denoting same-score comparison is annotated with the number of BTMs represented by the signature and all other annotations are the calculated Jaccard index. **(D)** Circos plot of BTM enrichment analysis across positive signature genes by viral family. Each sector represents a viral family, each point in all the tracks represents a BTM that was significant in at least one virus (padj <0.1). Track 2 is a barplot of the geometric mean of the expression of the genes represented by the BTM and plotted where the BTM was significant (padj <0.1). Each color in Track 3 is a granular annotation for each BTM pathway. The inner track connects the same BTM across viral families if they are both (left) positively or (right) negatively enriched. **(E)** Summary AUROC generated from the specific score (x-axis) across the different viral family dataset subsets (y-axis) comparing peak infection time category determined by 3B from healthy control animals. **(F)** AUROC of human data using the NHP Viral Response Signature (VRS) (n = 3183). **(G)** Violin plots of NHP VRS by viral severity of the samples from 3F. **(H)** Violin plots of NHP VRS by virus and disease of the samples from 3G. **(I)** Pearson’s correlation between the calculated MVS Score and the generated NHP VRS. Each dot is a single blood sample from an NHP across all timepoints collected (743 samples). **(J)** Pearson’s correlation between the calculated MVS (Human) score and the generated VRS (NHP) score. Each dot is a single blood sample from various public human gene expression datasets (n = 638). **(K)** Comparison of signature-enriched BTM pathways in the upregulated and downregulated genes in the MVS (Human) and VRS (NHP) pathways. Top 5 pathways (ordered by padj) chosen per signature’s up and down genes.

Next, we asked how conserved transcriptomic changes were across macaques between baseline and peak infection timepoints by viral family. We started by identifying gene signatures using our previously described multi-cohort analysis framework^17, 18^ to distinguish peak infection from the baseline T0 timepoint. We grouped the macaque datasets either by viral family – resulting in five signatures with one per viral family – or with all 21 datasets together to create the Viral Response Signature (VRS; Table S5). We compared each of these six newly developed signatures and the original MVS by gene with each other and by the blood transcription modules (BTM) that each gene set represented using the Jaccard index (JI; Figure 3C). While there was limited overlap between the virus-specific signatures at a gene-level (ranging from 3 shared genes between the Arenaviridae signature and Filoviridae signature to 32 overlapping genes between the Arenaviridae signature and Flaviviridae signature), there was much greater overlap in represented modules. Although some of the differences in genes may be due to variability in the statistical power, this result suggested that similar pathway networks are affected, even if driven by different genes (Figure 3C). Next, we performed enrichment analysis using the BTMs on the over- and under-expressed gene subsets from each signature separately (Table S6). We calculated a score for each BTM within the datasets grouped by viral families where the BTM was significant (padj<0.1; Figure 3D, track 2). Links connect BTMs across viral families where the BTM was enriched across the positive signature genes in both families. There were no connecting links across viruses by significant BTMs enriched in the negative signature gene subsets. Across all the viral families, there was a large number of upregulated pathways relating to myeloid and innate responses (Figure 3D). Similar analysis was also performed across all the DEGs identified at peak timepoints from Figure 3B and demonstrated similar results (Figure S7). We also assessed the generalizability of each virus-specific signature to other viral families. All virus-specific signatures distinguished healthy control and infected animals with high accuracy (AUROC>= 0.75; Figure 3E). By using this discovery/validation approach between viruses, this analysis demonstrates the robustly conserved innate responses that are upregulated across both hemorrhagic and nonhemorrhagic viral diseases.

We further investigated whether the VRS, identified using the macaque data, is applicable in distinguishing uninfected and infected human subjects. In 3183 human samples across 20 datasets of patients with one of 14 viral infections (Table S2), we found that the macaque VRS robustly distinguished viral infection from healthy across all symptomatic infections (Figures 3F and 3G). We separately looked at four human viral infections to demonstrate that the VRS signature robustly distinguished uninfected individuals from those infected with SARS-CoV-2, Ebola or dengue virus (padj<0.0001). Notably, the VRS was conserved upon infection by the Chikungunya virus (padj<0.0001), an RNA virus whose family was not included in the VRS signature discovery data (Figure 3H). Although VRS and MVS only had an overlap of 23 genes (JI = 0.03), these scores were significantly positively correlated across macaque data (r >= 0.54, p<1.4e-8) and human data (r >= 0.42, p<1.1e-8) across all timepoints collected, although they were noticeably lower for Flaviviridae viral infection in macaques (Figures 3I and 3J). In BTM overrepresentation analysis of the macaque-derived VRS signature compared to the human-derived MVS signature, upregulated genes in both signatures were enriched in innate response and antiviral modules, whereas downregulated BTMs corresponded to adaptive responses - further demonstrating conserved viral responses that transcend species and virus infections (Figure 3K and Table S7). Interestingly, while both signatures capture downregulation of genes associated with lymphoid cellular response modules, the negative genes represented in the VRS also captured non-lymphoid-specific responses such as signal transduction and translation initiation factor 3 (eIF3) pathways which may come from the inclusion of lethal, hemorrhagic viral diseases in macaque discovery datasets unlike MVS which was identified using respiratory viral infections. Together, these data validate macaques as robust models for studying human transcriptomic responses to viral infection. Here we demonstrate their potential for identifying both common antiviral responses as well as nuanced differences, such as those evident in Flaviviridae responses compared to other RNA viruses.

### Host response signature derived from macaques demonstrates robustness across acute and chronic viral infection in humans

Many patients have latent, chronic or acute viral infections that are not caused by single-strand RNA (ssRNA) viruses. However, the VRS and the MVS were identified using only acute infections caused by ssRNA viruses. Therefore, we investigated whether immune responses in macaques and humans were conserved across diverse viruses and disease manifestations. We used the macaque-derived VRS to further investigate the generalizability, and subsequently its translatability, to a variety of human viral infections.

First, across acute infections, the VRS score was significantly higher (p<0.05) in patients with Adenovirus (a double-stranded DNA (dsDNA) virus), Rotavirus (a double-stranded RNA (dsRNA) virus), Epstein-Barr virus (EBV; dsDNA virus) or human cytomegalovirus (HCMV; dsDNA virus) infections compared to healthy subjects (Figures 4A, 4B, 4C, and 4D). Second, we demonstrate that this response is robust in latent EBV (padj<0.01) but not in latent HCMV (padj=ns) infection (Figures 4C and 4D). Third, the VRS was also significantly higher (p < 0.01) in patients with chronic HIV (RNA virus with reverse transcription (RT) step), Hepatitis B (dsDNA virus with RT step), and Hepatitis C (ssRNA virus) viral infection (Figures 4E, 4F and 4G). Across the viruses studied here, the VRS demonstrated a stronger generalizability across represented viral infections in comparison to the MVS, potentially due to its discovery across a greater diversity of viral infections (Figure S8).

**Figure 4:**
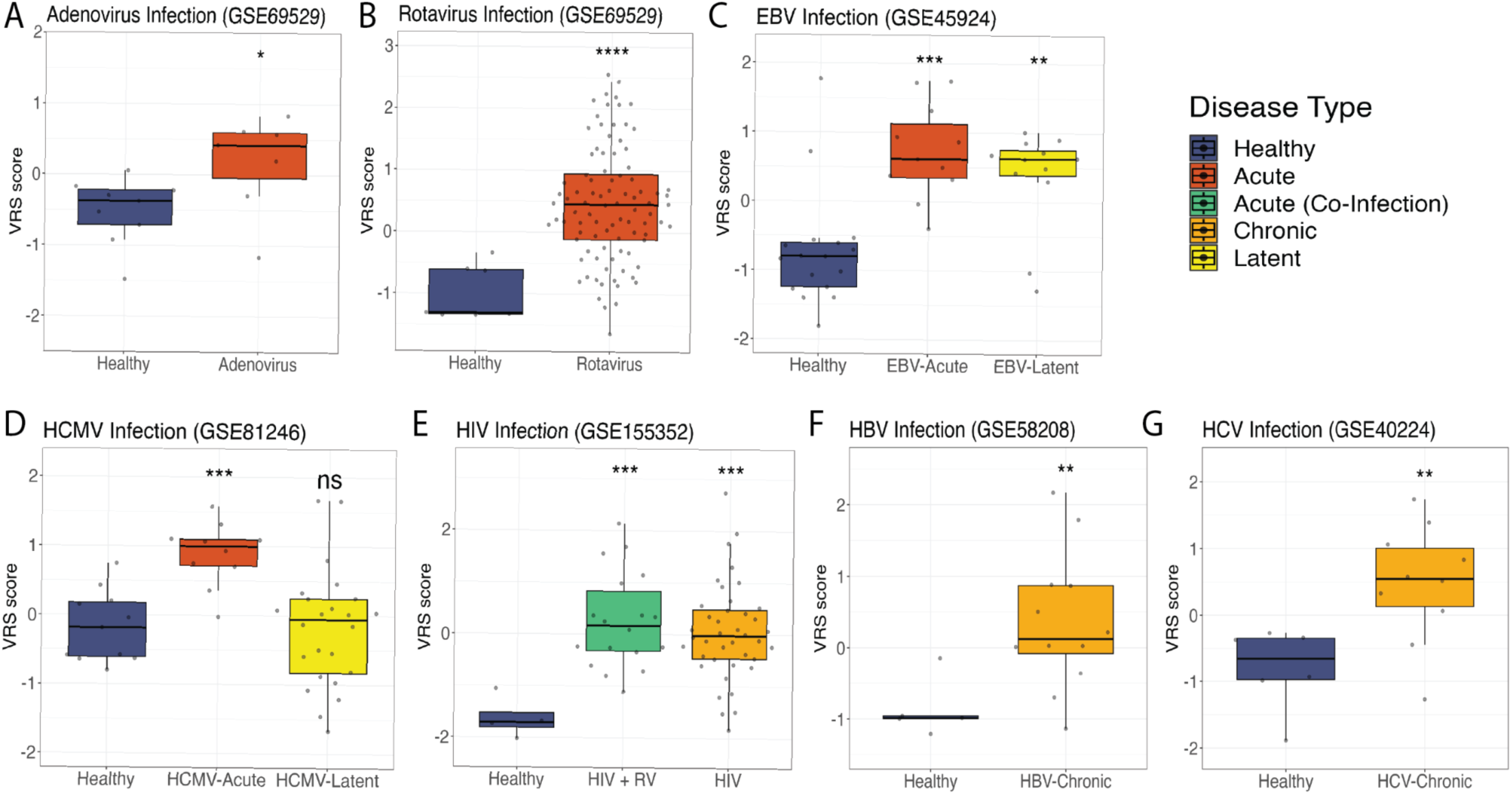
Macaque-discovered antiviral response is consistent in human acute and chronic, but not latent viral infections. **(A-G)** VRS score in blood samples from healthy control subjects versus patients with **(A)** Adenovirus infection, **(B)** Rotavirus infection, **(C)** acute or latent EBV infection, **(D)** acute or latent HCMV infection, **(E)** HIV infection or HIV co-infection with a respiratory virus (RV), **(F)** chronic HBV infection, and **(G)** chronic HCV infection. **(A-G)** Significance values were determined using an unpaired, one-sided Wilcoxon ranked-sum test looking at whether healthy VRS scores are less than comparator group VRS scores. Bonferroni correction for multiple hypothesis testing was applied per-subfigure and significance values were assigned by asterisk. Asterisk values across figure are represented as follows: *p value < 0.05, **p value < 0.01, ***p value < 0.001, and ****p value < 0.0001.

Consistently, the VRS distinguishes viral infection from a healthy state, and even detects chronic infection, regardless of viral genome strandedness or nucleic acid intermediates. This shows that a shared host response exists in both macaques and humans across various types of viral infections, highlighting the power of macaque models for understanding human antiviral responses.

### T cell responses differ across viral infections

Compared to other RNA viruses in macaques, the correlation between the VRS and MVS was noticeably lower, albeit significant, in Flaviviridae (Figure 3I and 3J). Both VRS and MVS signatures also had lower accuracy in distinguishing healthy controls and macaques with Flaviviridae infection. Therefore, we investigated potential drivers of this difference by evaluating the expression of four modules we have previously demonstrated to correlate with either severe or mild infection and be driven by different cell types.^8^ When comparing the expression of these four modules across datasets and at peak MVS score timepoints with the animals’ baseline samples, Module 4, composed of genes preferentially expressed in lymphoid cells (NK, T and B cells), was lower for all viral families except Flaviviridae in macaques (Figure 5A). In macaques infected with non-Flaviviridae viruses, Module 4 expression was inversely associated with the VRS score over time (Figure 5B and 5C). Module 4 was also significantly inversely correlated with VRS expression in humans infected with CHIKV, DENV, EBOV or SARS-CoV-2 (p<1.3e-5), though the correlation was lower in EBOV (r=-0.32, p=1.3e-5) and DENV infection (r=-0.47, p=2.3e-10; Figure 5D). We further looked at the association of Module 4 expression with viral disease severity across these human datasets and identified that while it significantly negatively associated with increased disease severity in SARS-CoV-2 and CHIKV infection (p<= 3.7e-10), the association was weaker in EBOV infection (p = 0.012) and not significantly associated with DENV disease severity (p = 0.12) (Figure 5E and 5F). Together, these data suggest a potential difference in lymphoid responses across different viral infections.

**Figure 5:**
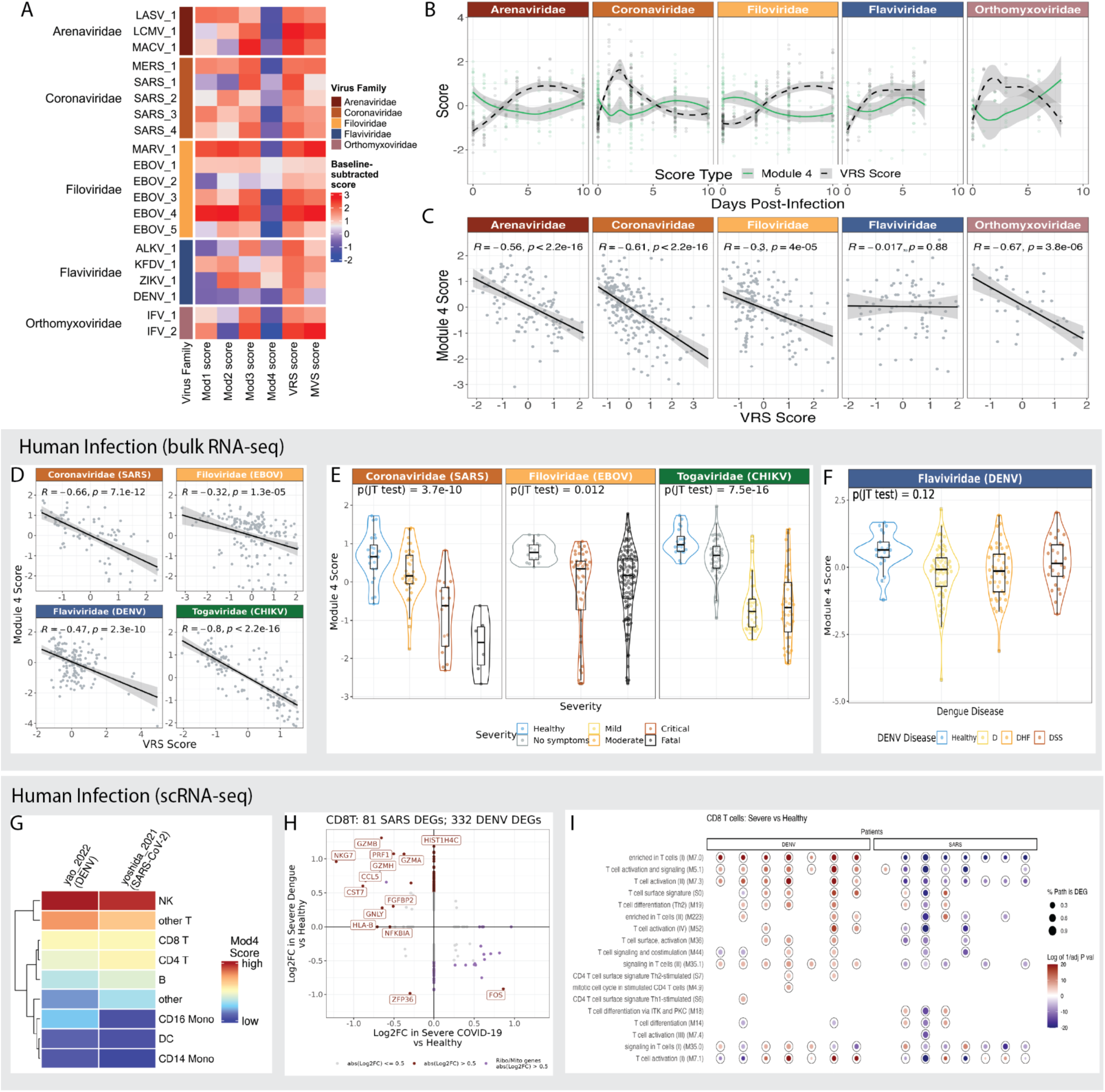
T cell responses differ between viruses in NHP and human viral infection. (A) Distribution of the Module 4 scores across macaques, comparing uninfected, healthy macaques to those at peak MVS score by viruses across 5 viral families. Each point represents a blood sample. Significance values were determined using an unpaired Wilcoxon ranked-sum test with Bonferroni correction for multiple hypothesis testing and assigned by asterisk. (B) Comparison of Module 4 scores to VRS scores across time in 4 viral families. (C) Comparison of Module 4 scores to VRS scores across the 4 viral species collected within the Flaviviridae family. (D) Module 4 score to VRS score in human data across 4 viral infections. (E) Module 4 scores by viral severity across CHIKV, EBOV, and SARS-CoV-2 viral infection. (F) Module 4 scores across disease timepoint and dengue disease type. (G) Expression of Module 4 scores by cell type in 2 scRNA-seq datasets. (H) Differential gene expression analysis of CD8 T cells across scRNA-seq data from COVID-19 and dengue patients between patients with severe disease compared to healthy controls. (I) BTM enrichment analysis of differentially expressed genes from each severe patient compared to the dataset’s healthy patients.

To understand what was driving a difference in lymphoid cellular responses, we identified two scRNA-seq datasets with healthy controls and patients presenting with severe disease from either SARS-CoV-2 or dengue viral infection (Figures S9A, S9B and S9C),.^19, 20^ We first investigated the cell types which had higher expression of Module 4 genes in patients with severe disease, and further identified the T cell and NK cell populations as drivers of these genes (Figure 4G and 4H). DEG analysis of these cell types demonstrated differences in the CD8 T cell responses between severe COVID-19 and dengue disease (Figure 5H; Table S8). In particular, genes marking effector and cytotoxic CD8 T cell profiles (*GZMB, NKG7, GZMH, PRF1, CCL5*) were upregulated in severe dengue and downregulated in severe COVID-19 (Figure 5H). Further, BTM enrichment using DEGs that were identified at a per virus-infected patient level in comparison to healthy controls showed that generally severe dengue patients upregulated genes related to T cell activation and differentiation modules whereas severe COVID-19 patients downregulated these module genes (Figure 5I). A similar trend was present in CD4 T cell responses (Figures S9E and S9F; Table S8), however, there were no strong differences in NK cell responses between severe COVID-19 and dengue disease (Figures S9G and S9H). This scRNA-seq analysis suggests a divergence in lymphoid responses between COVID-19 and DENV that may be linked to CD8 T effector functions and be important to address in vaccination strategies towards these different diseases.

## Discussion

Emerging and re-emerging viral diseases remain a constant global health threat mandating the development of new solutions to combat future epidemics and pandemics. Identification of conserved features of infection will enable the development of broad-spectrum antiviral solutions to fight constantly-evolving and emerging viruses. Multiple viruses of public health concern are understudied in humans; however, macaques remain an established model for understanding human disease with numerous independent virus studies published in these models providing a wealth of information on viral disease. Here, we reported that macaques are a reliable model of human antiviral transcriptomic responses by identifying and comparing conserved host responses across multiple viral families. We further extended the generalizability of conserved antiviral responses to several acute RNA viral infections of WHO priority concern, which include highly lethal viruses for which human data does not exist, and to viral infections by DNA viral infections and chronic viral diseases. Notably, we identified differences in longitudinal dynamics of antiviral responses and T cell functions depending on the infecting-virus. Together, these data provide detailed insights into the conserved, dynamic immune landscape of viral diseases across species that can provide potential targets for host-directed immunomodulatory antivirals.

We identified differences in the longitudinal dynamics of the conserved host response by virus across macaques and humans. These differences generally correspond with reported differences in incubation periods of these viruses in natural human disease: SARS-CoV-2, influenza, and HRV infection show shortened incubation periods relative to Ebola, Lassa and RSV.^21–23^ While different viral strains within the same viral species and other clinical variables have demonstrated effects on incubation periods (e.g., age, disease severity), our data indicates viral family level patterns as well.^24–26^ Additionally, even in longer-incubating viruses, our data demonstrated detectable induction of antiviral responses prior to peak infection that could potentiate development of viral diagnostics to identify patients prior to symptom presentation and/or viral shedding.

One of the reasons for differences in the longitudinal dynamics could be that viruses produce specific proteins to perturb host immune responses to support their undisturbed replication. While many viruses carry genes to produce anti-host proteins, differences in their cellular targets, number and efficacy may influence overall host antiviral dynamics. Another reason for these viral family-level differences may be variation in the ways these viruses present to pathogen recognition receptors (PRRs) (i.e., TLRs and RIG-I), which are important sensors for inducing antiviral immunity.^27^ Interestingly, Arenaviridae, Filoviridae (Order: *Mononegavirales*), and RSV (Order: *Mononegavirales*) are negative-stranded RNA viruses (-ssRNA) that replicate in the cytoplasm, differing from the nucleus-replicating -ssRNA influenza virus and positive-stranded RNA viruses (+ssRNA) such as Coronaviridae and Flaviviridae viruses. This difference in genome strandness may be important as studies have shown that during replication, unlike +ssRNA, dsRNA and DNA viruses, -ssRNA viruses tend to produce low levels of dsRNA intermediates – robust triggers of antiviral responses capable of binding directly to RIG-1 and MDA-5 receptors for activation of IRFs and NF-kB for downstream interferon and pro-inflammatory responses.^28–30^ Additionally, -ssRNA viruses cannot be directly translated but need to generate +ssRNA to co-opt host translation machinery to reproduce, which may delay their overall infection dynamics. While it has been posited that other proteins and complexes produced by -ssRNA can drive inflammatory responses,^31^ differences in timing of protein production based on starting nucleic acid material within cells could also drive differences in overall timing of the induction of host antiviral responses. Other factors related to the viral family-level differences could be the differences in route of infection, viral tissue tropism, and viral latency. For example, influenza viral shedding (via nasal wash) is generally detectable via PCR one day after viral challenge and before symptom onset, whereas EBOV is generally detected after symptom onset in the blood.^32, 33^ While our study focuses on cells in the blood, there could be differences in the types of tissues infected and the magnitude of cytokine and ISG production, resident cell activation, and immune cell recruitment detectable in peripheral blood. Further studies on comparative viral immunology and disease are required to ascertain the drivers of these different dynamics. However, as it stands this data suggest key post-infection timepoints for study design and sample collection based on the infecting viral family, which could be useful to development of macaque models for new viruses.

Here we report that a conserved set of genes identified across viral infection of macaques is robust across infection of different macaque species and heterogenous human populations by viruses across the Baltimore Virus Classification system. While focused studies within viral families have compared transcriptional responses between different macaque species to suggest conservation in transcriptional responses - ie SARS-CoV-2 infection between cynomolgus and rhesus macaques, SARS-CoV-2 infection between green monkeys and rhesus macaques, and EBOV infection between rhesus and cynomolgus macaques – this study is the first that we are aware of that directly translates NHP findings to humans.^34–36^ This step is important because NHP studies allow for easier design of randomized control trials and challenge studies testing various interventions (therapeutics, vaccinations, infection strains, etc.) and this work suggests measuring a readout in the form of changes to the VRS could translate to human-relevant interpretations. Additionally, here we are showing that these responses are conserved in acute infection by pathogens across six out of seven Baltimore Classification groups (we could not test any ssDNA viruses because of the lack of available public data). This further highlights that despite the diverse genetic backbones and intermediates prior to mRNA production, these viruses still are detectable by the host to drive similar antiviral responses. This is not unexpected: recent work suggests that while DNA and RNA sensors rely on different mechanisms across various cellular compartments, there is appreciable cross-talk between these different pathways.^37^ For example, the cGAS-STING axis, established as a cytosolic DNA sensing pathway, has been shown to be involved in responses to RNA viruses including dengue virus, vesicular stomatitis virus and influenza A virus with many mechanisms hypothesized, including STING binding to RIG-I/MAVS (cytosolic RNA sensors) that carries out signaling downstream of their activation.^38–43^ These data further suggest that while virus-specific gene sets identified may not consist of precisely the same genes, they are part of similar pathways, indicative of redundancy in the antiviral response that may be broadly targetable.^44, 45^ These differences may also be driven by virus-specific host targets that could induce specific gene expression changes within pathways.

While many macaque models of RNA viral infections demonstrate clinical presentation, viremia, and virus tissue tropism that is similar to their respective human viral disease; there exist multiple viruses that show divergence in pathogenesis between primate species.^46–48^ For example, rhesus macaques infected with COVID-19, dengue or Kyasanur Forest Disease virus (KFDV) have viremia, yet do not recapitulate human disease phenotypes.^46, 47, 49^ Even so, we show that both the MVS and VRS can still detect macaque infection in cases of COVID-19 or dengue viral infection where macaques are lacking in clinical symptoms (asymptomatic or mild disease). An important divergence between rhesus macaques and human genotypes that may account for differences in responses may be driven by the amino acid composition of the STING protein, which plays a role in antiviral responses through IRF3 activation and downstream ISG expression. Thus, while DENV can cleave human STING to reduce type I IFN production and enhance viral replication, it cannot cleave rhesus macaque STING and restrict antiviral pathways downstream of STING activation. This evidence aligns with the theory that rhesus macaques form part of dengue virus’ sylvatic cycle, asymptomatically maintaining the virus population in nature. Even across macaque species, while rhesus, pigtail, and cynomolgus macaques are highly genetically similar with comparable susceptibility to many of the same viral infections, there exist nuances in their genotypes that make some macaque species better models of human viral disease than others. In particular, tripartite motif-containing protein 5 (TRIM5) is an interferon-induced (IFI) antiviral protein that has multiple isoforms. While rhesus macaques express heterogeneous TRIM5 genotypes, pigtail macaques only express one genotype for this protein. In infection by simian immunodeficiency virus (SIV), rhesus macaques’ TRIM5 can differentially restrict SIV unlike pigtail macaques’ TRIM5, thereby leaving pigtail macaques susceptible to SIV infection, making them the preferred NHP model for studying HIV. TRIM5 is also important for restricting the NS2B/3 protein of KFDV and other tick-borne flaviviruses to limit their replication.^50, 51^ It has been hypothesized that the specific TRIM5 protein expressed by pigtail macaques reduces this macaque’s ability to inhibit KFDV NS2B/3, thereby making pigtail macaques more susceptible to and clinically symptomatic upon KFDV infection as compared to rhesus macaques which present with asymptomatic and mild Kyasanur Forest disease.^50, 51^ Differences in pathogenicity between macaques and humans may suggest targets for antivirals, such as blocking DENV’s cleavage of STING or introducing enzymes to optimally degrade flavivirus NS2B/3. Together, these data demonstrate that even with genotype differences that may impact the translation of results from macaque to humans (i.e. where STING and TRIM5 pathways are involved), the MVS and VRS magnitudes may provide a proxy for disease detection and intervention assessment even when clinical symptoms are not present.

While T cells are known to be crucial for mounting effective immune responses during viral infections, we found opposite trends in T cell transcriptional responses between dengue and COVID-19 during severe disease manifestations. During COVID-19, lymphopenia and exhausted T cells are found in patients with severe and fatal disease outcomes and may serve as a potential prognostic for disease outcome.^52, 53^ Lymphopenia has also been described in severe Ebola infections while exhausted T cells have also been a marker of chronic viral infections such as HIV, Hepatitis B, and Hepatitis C.^54, 55^ In contrast, the role of T cell responses in dengue has been highly disputed. The majority of symptomatic dengue disease is driven by secondary dengue infections, yet it is unclear whether pre-existing dengue-specific T cells play a protective or pathogenic role. Generally, studies consistently report an expansion of pre-existing T cell populations readily activated upon secondary dengue infection^56^, which is in line with our data. However, while some studies report no transcriptional differences in the quantity and quality of DENV-specific CD4 and CD8 T cell populations by disease severity,^57, 58^ others suggest that the expansion of pre-existing cross-reactive T cells drives severity through ineffective viral control and aberrant cytokine responses.^59–61^ Though we cannot distinguish DENV specificity or pathologic versus protective functional differences, we do see a general activation of the T cell compartment. Interestingly, all macaques had a primary flavivirus infection and consistently demonstrated differing trends in the expression of MVS genes relating to B, T and NK cell responses (Module 4) as compared to other viral infections. This result suggests Flaviviridae-specific differences in T cell response induction independent of prior exposure. These data together are important because they highlight general T cell activation and proliferation may not directly equate to protective viral responses, and suggest further work needed in dissecting specific T cell subsets and their role across viral disease pathogenicity.

In conclusion, we conducted a comprehensive analysis of host responses to a wide range of viruses, using transcriptomic data from both human and macaque cohorts. Leveraging macaque data uniquely allowed us to gather pre-symptomatic, post-inoculation timepoints from infection by pathogenic and lethal viruses that are otherwise very difficult, if not impossible, to obtain from humans. Our integrated analysis across heterogeneous macaque and human cohorts allowed us to identify highly generalizable antiviral responses conserved in acute and chronic viral disease. Moreover, our analyses identified differences in the longitudinal dynamics of host response induction and resolution, which appear to be influenced by viral family. These results further support the reliability of macaque models in studying human antiviral responses and are useful for pandemic preparedness. Specifically, our work identifies several key areas for future research and development of antiviral countermeasures, including the design of new intervention strategies such as diagnostics and therapeutic timing, the optimization of macaque challenge study design for emerging and reemerging viruses, and the development of broad-spectrum host-directed immunomodulatory therapies. These findings underscore the importance of continued comparative research across transcriptomic responses to diverse viruses in macaques and humans, with the ultimate goal of improving our ability to predict, prevent, and treat viral infections.

### Limitations of Study

Our study has a few potential limitations. First, in a number of microarray datasets, not all genes in each gene set tested were present, in which case we measured the subset of genes that were present in the particular dataset. Our previous work has shown that many of the genes within the MVS score are highly correlated and only a subset is required to detect conserved responses. Second, all macaque genes were converted to human homologues, ignoring the expression of macaque genes that do not have a clear human homologue and/or may be important to overall antiviral response dynamics. Third, we only analyzed transcriptome data from blood samples. However, differences in antiviral responses may occur at the tissue level and site of infection that we may have missed. Fourth, some viral families we analyzed only had a small representation of viral species and did not have different viral variants accounted for. While we use grouping at the viral family level to identify conserved patterns, these may not be applicable to every virus within that viral family. For example, not all viruses cause symptomatic disease in both macaques and humans, such as SIV in humans and HIV in macaques. While we included all macaque viral datasets possible and a broad range of viruses with epidemic and pandemic concern, MVS and VRS responses need to continually be tested in new viral infection datasets.

## STAR★Methods

### Key Resources Table

| REAGENT or RESOURCE | SOURCE | IDENTIFIER |
| --- | --- | --- |
| **Deposited data** |  |  |
| Microarray dataset | Fukuyama et al., 2020 | accession# GSE152406 |
| Microarray dataset | Skinner et al., 2014 | accession# GSE60009 |
| Microarray dataset | Malhotra et al., 2013 | accession# GSE41752 |
| Microarray dataset | Rasmussen et al., 2014 | accession# GSE49838 |
| Microarray dataset | Djavani et al., 2007 | accession# GSE5790 |
| Microarray dataset | Connor et al., 2015 | accession# GSE58287 |
| RNA-seq dataset | Speranza et al., 2018 | accession# GSE103825 |
| Microarray dataset | Yen et al., 2010 | accession# GSE24943 |
| RNA-seq dataset | Maroney et al., 2021 | accession# PRJNA718880 |
| RNA-seq dataset | Versteeg et al., 2017 | accession# PRJNA398558 |
| RNA-seq dataset | Speranza et al., 2017 | accession# GSE99463 |
| RNA-seq dataset | Broeckel et al., 2021 | accession# GSE185797 |
| Microarray dataset | Aid et al., 2017 | accession# GSE90868 |
| Microarray dataset | Strouts et al., 2016 | accession# GSE72430 |
| Microarray dataset | de Wit et al., 2013 | accession# GSE44542 |
| RNA-seq dataset | Aid et al., 2020 | accession# GSE156701 |
| RNA-seq dataset | Price et a., 2020 | accession# GSE155363 |
| RNA-seq dataset | Coleman et al., 2021 | accession# GSE184949_GPL29319 |
| Microarray dataset | Rubins et al., 2006 | accession# GSE4013_GPL3093 |
| Microarray dataset | Rubins et al., 2006 | accession# GSE4013_GPL3346 |
| Microarray dataset | Rubins et al., 2006 | accession# GSE4013_GPL3347 |
| RNA-seq dataset | Reynard et al., 2023 | accession# GSE158390 |
| scRNA-seq dataset | Kotliar et al., 2020 | Zenodo# 7229439 |
| RNA-seq dataset | Liu et al., 2017 | accession# PRJNA352396 |
| RNA-seq dataset | Soares-Schanoski et al., 2019 | accession# PRJNA507472 |
| RNA-seq dataset | Michlmayr et al., 2018 | accession# PRJNA390289 |
| RNA-seq dataset | Thair et al., 2020 | accession# GSE152641 |
| Microarray dataset | de Steenhuijsen Piters et al., 2016 | accession# GSE77087 |
| Microarray dataset | Liu et al., 2016 | accession# GSE73072 |
| Microarray dataset | Zhai et al., 2015 | accession# GSE68310 |
| Microarray dataset | Jaggi et al., 2018 | accession# GSE68004 |
| Microarray dataset | Heinonen et al., 2016 | accession# GSE67059 |
| Microarray dataset | Sweeney et al., 2015 | accession# GSE66099 |
| Microarray dataset | Wong et al., 2007, Wong et al., 2008, Wong et al., 2012 | accession# GSE4607 |
| Microarray dataset | Ramilo et al., 2007 | accession# GSE6269 |
| Microarray dataset | Hoang et al., 2014 | accession# GSE61821 |
| Microarray dataset | Davenport et al., 2014 | accession# GSE61754 |
| Microarray dataset | Parnell et al., 2012 | accession# GSE40012 |
| Microarray dataset | Mejias et al., 2013 | accession# GSE38900 |
| Microarray dataset | Wang et al., 2007 | accession# GSE2729 |
| Microarray dataset | Berdal et al., 2011 | accession# GSE27131 |
| Microarray dataset | Smith et al., 2014 | accession# GSE25504 |
| Microarray dataset | Bermejo-Martin et al., 2010 | accession# GSE21802 |
| Microarray dataset | Parnell et al., 2011 | accession# GSE20346 |
| Microarray dataset | Zaas et al., 2009 | accession# GSE17156 |
| Microarray dataset | Yu et al., 2019 | accession# GSE117827 |
| Microarray dataset | Dunning et al., 2018 | accession# GSE111368 |
| Microarray dataset | Rodriguez-Fernandez et al., 2017 | accession# GSE103842 |
| Microarray dataset | Tang et al., 2019 | accession# GSE101702 |
| Microarray dataset | Jong et al., 2016 | accession# E-MTAB-5195 |
| Microarray dataset | Simmons CP et al. | accession# GSE40628 |
| Microarray dataset | Long HT et al. | accession# GSE13052 |
| Microarray dataset | Popper SJ et al. | accession# GSE38246 |
| Microarray dataset | Kwissa M et al. | accession# GSE51808 |
| RNA-seq dataset | Daberg et al. | accession# GSE69529 |
| RNA-seq dataset | Sellers et al. | accession# GSE155352 |
| Microarray dataset | N/A | accession# GSE58208 |
| Microarray dataset | Dunmire et al. | accession# GSE45924 |
| Microarray dataset | Riou et al. | accession# GSE81246 |
| Microarray dataset | Bolen et al. | accession# GSE40224 |
| **Software and Algorithms** |  |  |
| R | R Core Team (2022) | https://www.r-project.org/ |
| MetaIntegrator | Haynes et al. | https://cran.r-project.org/web/packages/MetaIntegrator/index.html |
| Seurat | Satija et al., 2015 | https://satijalab.org/seurat/ |
| DESeq2 | Love et al., 2014 | https://bioconductor.org/packages/release/bioc/html/DESeq2.html |
| ComplexHeatmap | Gu, 2022 | https://bioconductor.org/packages/release/bioc/html/ComplexHeatmap.html |
| lmerTest | Kuznetsova et al., 2017 | https://cran.r-project.org/web/packages/lmerTest/index.html |
| scanpy | Wolf et al., 2018 | https://scanpy.readthedocs.io/en/stable/ |
| ggplot2 | Wickham H, 2016 | https://ggplot2.tidyverse.org/ |
| UMAP | McInnes et al., 2018 | https://cran.r-project.org/web/packages/umap/index.html |
| Tximport | Soneson and Robinson, 2018 | https://bioconductor.org/packages/release/bioc/html/tximport.html |
| Salmon | Patro et al., 2017 | https://combine-lab.github.io/salmon/ |

### Resource Availability

#### Lead Contact

Further information and requests for resources, software, and data should be directed to and will be fulfilled by the Lead Contacts, Catherine Blish and Purvesh Khatri.

#### Materials Availability

This study did not generate new unique reagents.

#### Data and Code Availability

This study did not generate any unique datasets or code. All datasets, software, and algorithms used in this study are publicly available and listed in the Key Resource table. Code used to generate figures also available on Github: https://github.com/Khatri-Lab/NHP_virus_challenge.

#### Method details

Methods for analyses performed are described below.

### Quantification and Statistical Analysis

#### Non-human primate dataset collection and preprocessing

We downloaded 21 gene expression datasets (either microarray or RNA-seq) from the National Center for Biotechnology Information (NCBI) Gene Expression Omnibus (GEO), Sequence Read Archive (SRA) or shared by collaborators, consisting of 743 samples derived from whole blood or peripheral blood mononuclear cells (PBMCs) (Table 1, Table S1). The counts dataset generated by Reynard et al.^11^ was downloaded from Zenodo. The samples in these datasets included all available macaque virus challenge studies with samples from uninfected and two or more infected timepoints. We incorporated technical heterogeneity in our analysis as these datasets were profiled using microarray and RNA sequencing (RNA-seq) from different manufacturers. Raw RNA-seq reads were trimmed of Illumina adaptors and reads that were too short after adaptor trimming (less than 20 nt) were removed using Trim Galore (v0.6.5). We then mapped the cleaned reads to the macaque transcriptome (Salmon v1.3.0, genome version Mmul_10 or Macaca_fascicularis_6.0).^62^ We used Tximport (v1.26.1)^63^ to summarize to gene-level expression. Finally, we applied the variance stabilizing transformation from DESeq2 (v1.38.2)^64^ to normalize gene expression for downstream analysis and visualization. Within a dataset, cohorts assayed with different microarray types or had different viruses with independent control animals were treated as independent.

We mapped all genes generated through alignment to macaque genomes (rhesus: Mmul_10, cynomolgus: Macaca_fascicularis_6.0, pig-tailed: Mnem_1.0) to the corresponding human orthologs to facilitate integrated, comparative analyses. For the scRNA-seq data, Kotliar et al.^16^ generously shared their scRNA-seq object of GSE158390 that was processed and annotated for cell type.

#### Human dataset collection and preprocessing

We utilized human gene expression data collected and processed across our other studies ^8, 65, 66^. Briefly, 47 gene expression datasets from the National Center for Biotechnology Information (NCBI) Gene Expression Omnibus (GEO), Sequence Read Archive (SRA), ArrayExpress, and European Nucleotide Archive (ENA), consisting of 5345 samples derived from whole blood or peripheral blood mononuclear cells (PBMCs) (Table S2). Log2 transformation and quantile normalization was applied when necessary. For the combined discovery dataset from Zheng et al.^8^ (Table S2) and the human dengue datasets,^67–70^ Combat CONormalization Using conTrols (COCONUT) was used for between-dataset normalization.^71^ Healthy samples from each cohort undergo ComBat co-normalization without covariates, and the ComBat estimated parameters are computed for the healthy samples in each dataset. By applying these parameters to the non-healthy samples, all datasets keep the same background distribution while retaining the same relative distance between healthy and disease samples, which preserves the biological variability between the two groups within a dataset.

#### Gene signatures and scoring

We used a number of previously published gene signatures, including: MVS^7^, MS1 signature^10^, T cell activation signature^72^, and ISG signature^73^. We generated gene signatures across all macaque datasets, the RNA virus subset of the datasets, and each viral family subset of the datasets using the MetaIntegrator workflow. Briefly, score generation was done by applying two meta-analysis methods previously described: (1) combining effect sizes and (2) combining p values. To generate robust, comparable gene signatures per virus, we filtered signatures for effect size of 0.6 and false discovery rate (FDR) thresholds between 0.05 to 0.0001 in order to find thresholds that captured around 200 genes for better comparison across gene signatures. For the Viral Response Signature (VRS) that was generated across all macaque datasets, we removed one dataset at a time and applied both meta-analysis methods at each iteration to avoid the influence of any datasets with large sample sizes on the results.

We defined each score by the geometric mean of the normalized, log2-transformed expression of the overexpressed genes minus the geometric mean of the normalized, log2-transformed expression of the underexpressed genes of each gene signature. We scaled and centered (mean = 1, standard deviation = 1) all sample scores per dataset to allow for comparison between datasets.

We measured the correlation between different scores using Spearman’s rank correlation coefficient. We used the Mann– Whitney U test (Wilcoxon rank-sum test) to compare MVS scores between two groups.

#### Mixed-effects Model for Timepoint Data

We used multivariable linear mixed-effects models with random time-influenced subject-specific intercepts and slopes to assess the changes in MVS scores from uninfected baseline timepoints (intercepts), and follow-up timepoints post-infection (slopes). Separate models for were estimated using time, day-post-infection * day-post-infection, macaque species, dataset, and infecting viruses that included various interactions between these covariates. Across these various models, the one we chose was that with the lowest Akaike Information Criterion (AIC) value. The final reported model for both the macaque and the human datasets was: lmer formula = MVS_score ∼ Time + Time^2^ + Virus + Time*Virus_Family + Time^2^*Virus_Family+(1+Time|Subject). Analyses were run in R version 4.2.2 using the “lmerTest” package.

#### Gene Set Overrepresentation Enrichment Analysis

Overrepresentation analysis was performed on differentially expressed gene sets or signature sets identified from bulk RNA-seq (padj < 0.05 and ES >= 0.1) and/or scRNA-seq (padj < 0.05 and ES >= 0.6) analyses utilizing the Blood Transcriptional Modules (BTM). BTMs for which there was higher level annotation from Hagan et al^74^ were visualized in Circos plots and single cell analysis. The p values were adjusted using Bonferroni correction. Data analysis was performed using R.

#### Analysis of single-cell RNA sequencing

Data from Kotliar et al.^16^ was generously shared as a processed object with cell types already assigned. We generated gene scores per cell type utilizing the geometric mean of the genes in the signature. Processed data from Yoshida et al.^20^ was downloaded from GEO. Data from Ghita et al.^19^ was generously shared. Both datasets were processed via Seurat and scanpy for QC, dimension reduction, clustering, and cell type classification. Seurat v4^75^ was used for cell type annotation utilizing the multimodal PBMC reference dataset from the associated publication, and cell type calls were compared to previous manual annotation of datasets for confirmation. We generated MVS^7^ and Module 4^8^ scores per celltype across both datasets using the geometric mean of the genes in the subset. We utilized the FindMarkers function in Seurat to perform DEG analysis on each viral infected individual compared with all healthy controls in each dataset by cell type. We then performed BTM enrichment analyses per individual on the gene subset that was upregulated upon infection and the gene subset that was downregulated upon infection separately (padj < 0.05 and ES >= 0.6). The pathway direction that had the highest adjusted p- value was retained if it appeared in both the up and down regulated module list.

#### Figure Generation

Figures were generated in R using the “ggplot2*”* and ComplexHeatmap package. Colors for figures were generated using the “NatParksPalettes*”*package. Statistical analyses were performed as described in figure and table legends and plotted using the R “ggpubr’ package.

## ACKNOWLEDGMENTS

We are grateful to all the labs that conducted the experiments and generated and publicly shared their data. We also thank all study participants and their families for being involved in all included studies. We thank Dylan Kotliar for sharing processed scRNA-seq data with us. Finally, we thank the members of the Blish Lab and Khatri Lab for useful suggestions: Denis Dermadi, Maïgane Diop, Mike Freedman, Ananthakrishnan Ganesan, Umay Geyikci, Sanjana Gupta, Rebecca Hamlin, Larry Kalesinskas, Ian Lee, Maddie Lee, Michelle Leong, Yiran Liu, Giovanny Martinez-Colón, Andrew Moore, Ruoxi Pi, Kass Pinedo, Thanmayi Ranganath, Izumi de los Rios Kobrara, Makeda Robinson, Sonieda Rodriguez, Arjun Rustagi, Sarah Sackey, Marion Santo, Ben Solomon, Mikayla Stabile, Simone Thair, Aaron Wilk, and Mengyang Zhang.

K.R. is funded by the National Science Foundation Graduate Research Fellowship 2019282939 and Bio-X graduate Stanford Graduate Fellowship. J.T. is funded by National Science Scholarship (PhD) from the Agency of Science, Technology, and Research (A∗STAR), Singapore. Z.Y. was supported by a Thrasher Research Fund early career award program grant and by a postdoctoral fellowship from the Maternal and Child Health Research Institute, Lucile Packard Foundation for Children’s Health. V.D. was supported by a Chan Zuckerberg Biohub Collaborative Postdoctoral Fellowship. S.E. is funded by a NIAID grant RO1AI158569, an Investigator Initiated Award number W81XWH2210283 and an expansion award number W81XWH2110456 from the DoD office of the CDMRP/PRMRP, and a Defense Threat Reduction Fundamental Research to Counter Weapons of Mass Destruction grant HDTRA11810039. S.E. and C.A.B. are Investigators of the Chan Zuckerberg Biohub. C.A.B. is funded in part by NIH DP1 DA046089 (C. A. B.), a 2019 Sentinel Pilot Project from the Bill and Melinda Gates Foundation and OP113682 and U19AI057229 from the Bill and Melinda Gates Foundation. P.K. is funded in part by the Bill and Melinda Gates Foundation (OPP1113682); the National Institute of Allergy and Infectious Diseases (NIAID) grants 1U19AI109662, U19AI057229, and 5R01AI125197; Department of Defense contract W81XWH-18-1-0253. S.E. and P.K. are funded in part by Department of Defense contract W81XWH1910235 and the Ralph & Marian Falk Medical Research Trust. The funders had no role in study design, data collection and analysis, decision to publish, or preparation of the manuscript.

## AUTHOR CONTRIBUTIONS

K.R, C.A.B and P.K. conceived the study. C.A.B and P.K. supervised the study. K.R., H.Z., J.T. and M.D. collected, annotated, processed, and analyzed data. K.R., C.A.B and P.K. interpreted analysis results and wrote the manuscript. S.E. supervised the dengue scRNA-seq study cohort. V.D. and S.E. designed study and enrolled patients with dengue infection. Z.Y. profiled and processed single-cell RNA-seq of PBMCs from patients with dengue infection. M.K., M.R., J.M.T., M.G., K.E.F., J.R.F., K.S.C., D.C.D., and R.A.S. designed and carried out the study profiling RNA-seq of PBMCs from NHPs with SARS-CoV-2 infection.

## SUPPLEMENTAL FIGURES/ TABLES

**SFig1:**
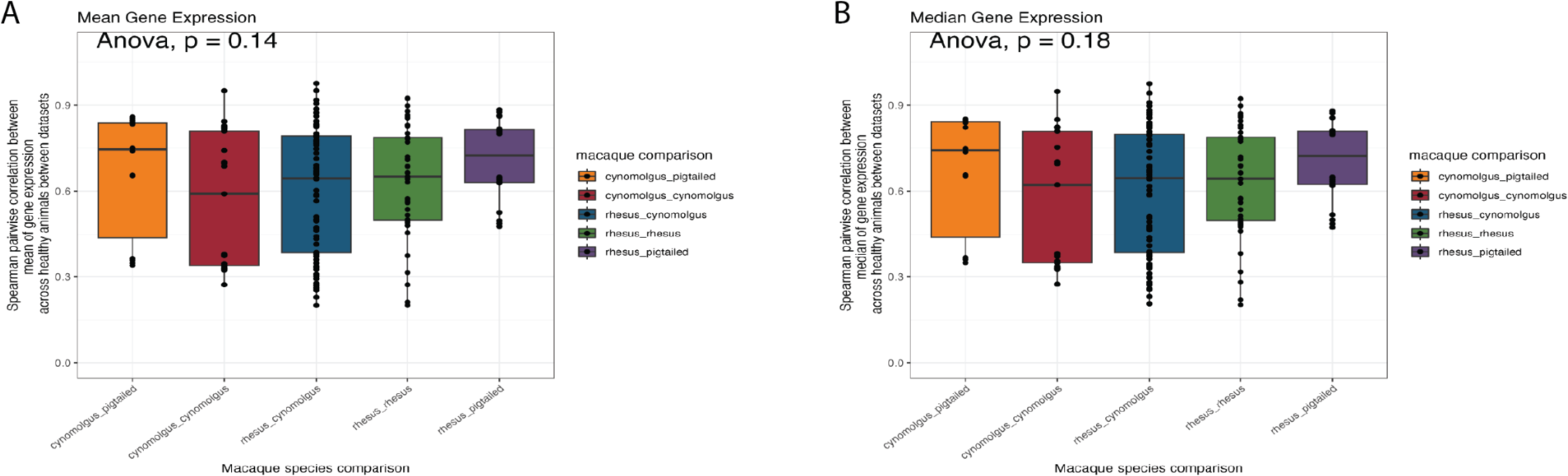
Gene pair correlation comparison across the macaques. (A-B) Correlation coefficients between the (A) mean and (B) median gene expression of healthy animals per dataset in comparison to other comparison datasets from the same or other macaque species.

**SFig2.**
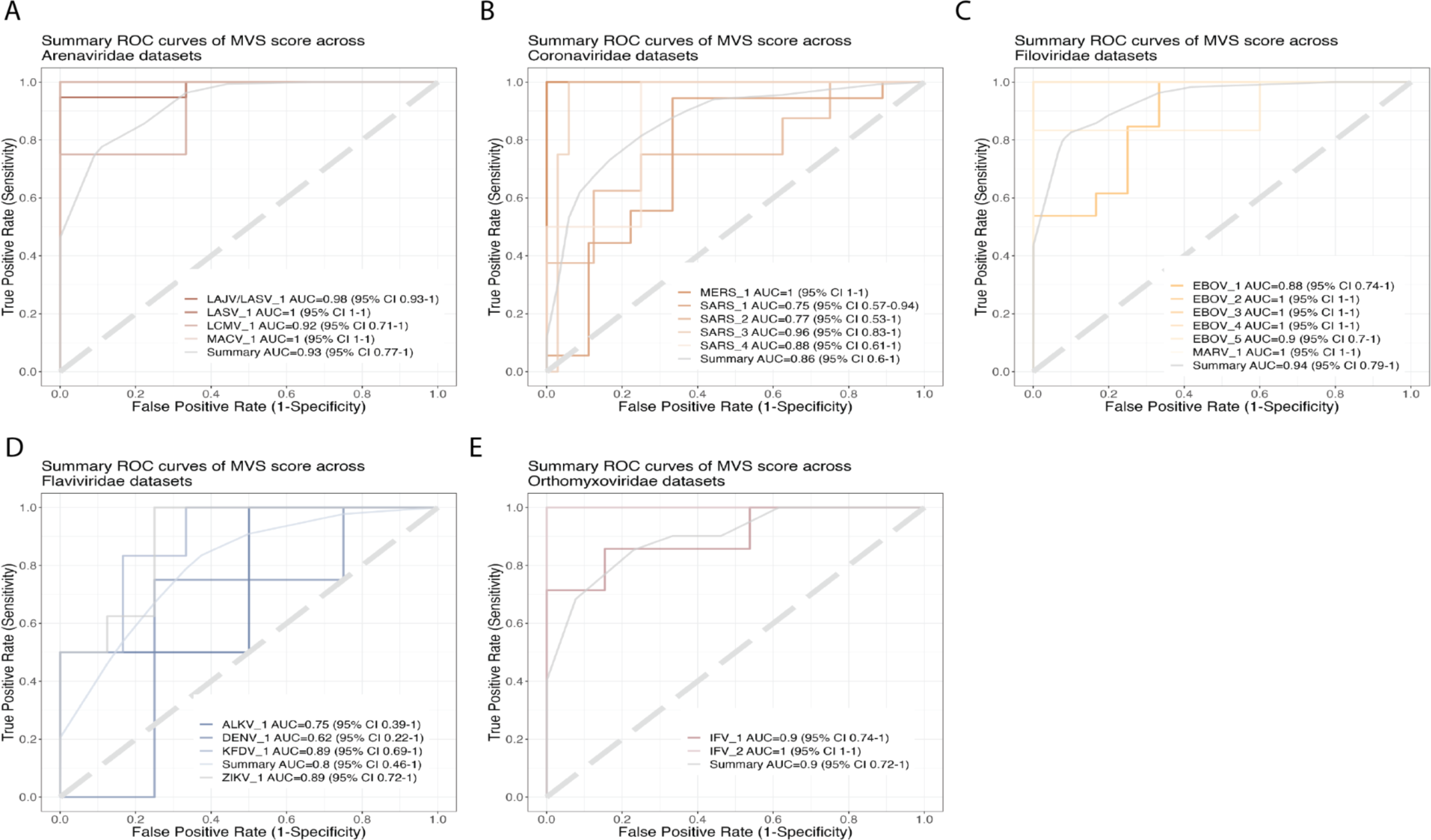
AUROCs by dataset. (A-E) ROC curves for distinguishing macaques with viral infection at peak MVS timepoint category from uninfected macaques, across datasets by viral family and colored by individual dataset.

**SFig3.**
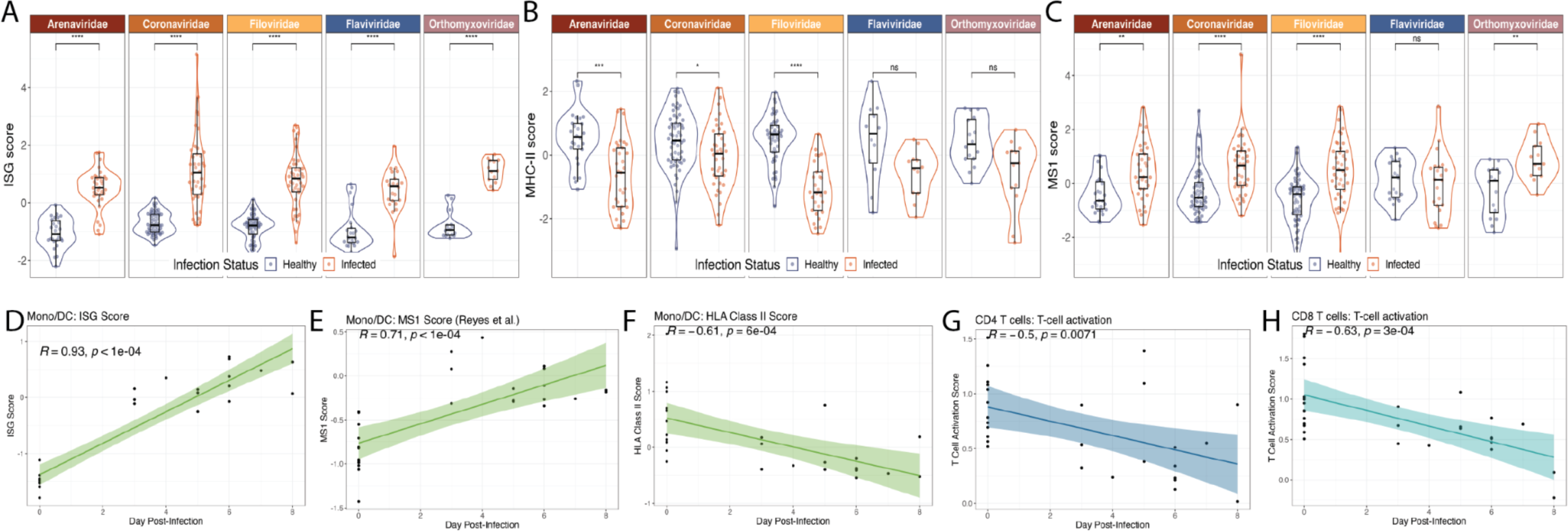
Validation of independent inflammation and monocyte and T cell function scores in NHP data. (A-E) Gene score by sample taking the geometric mean of the genes present in the scaled (A) ISG score, (B) HLA Class II Score, (C) MS1 Score, (D-F) Correlation of ISG, HLA class II and MS1 scores with time post-infection, (G-H) T cell activation score and plotting the average by macaque and timepoint.

**SFig4.**
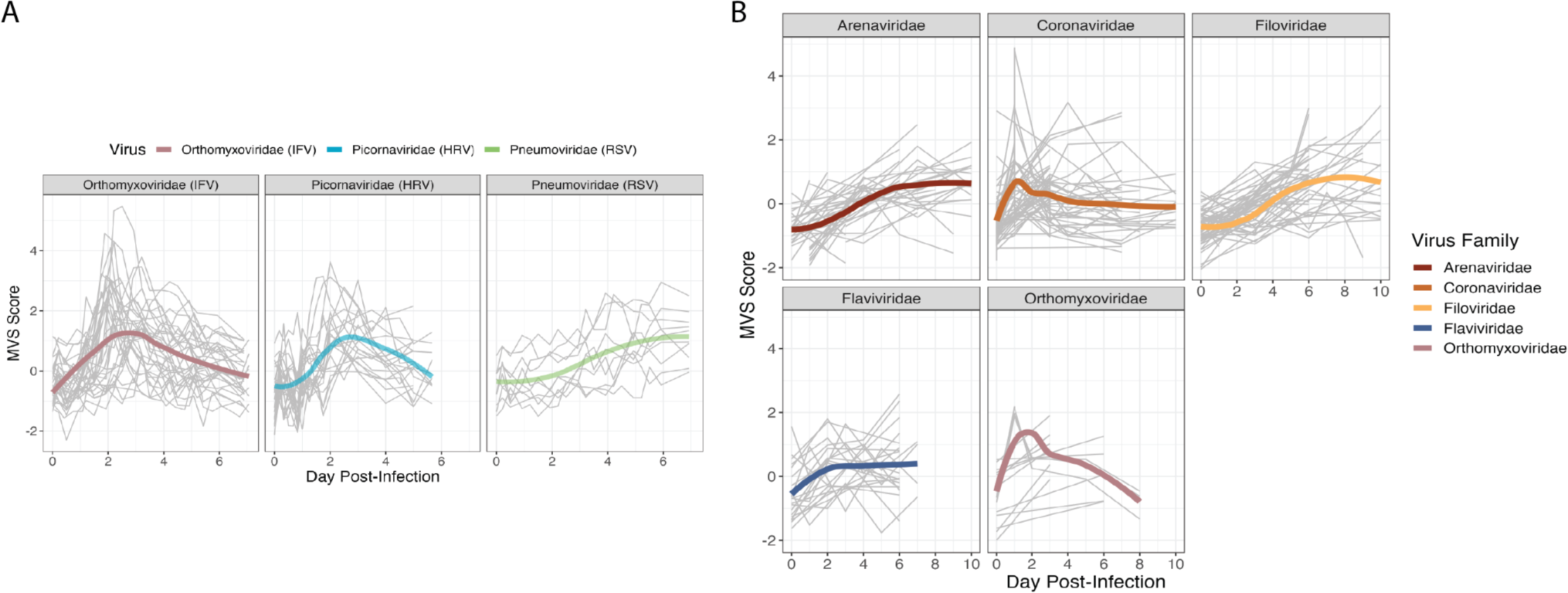
MVS score across all data timepoints. (A-B) MVS score calculated across (A) human longitudinal datasets and (B) all NHP longitudinal data.

**SFig5.**
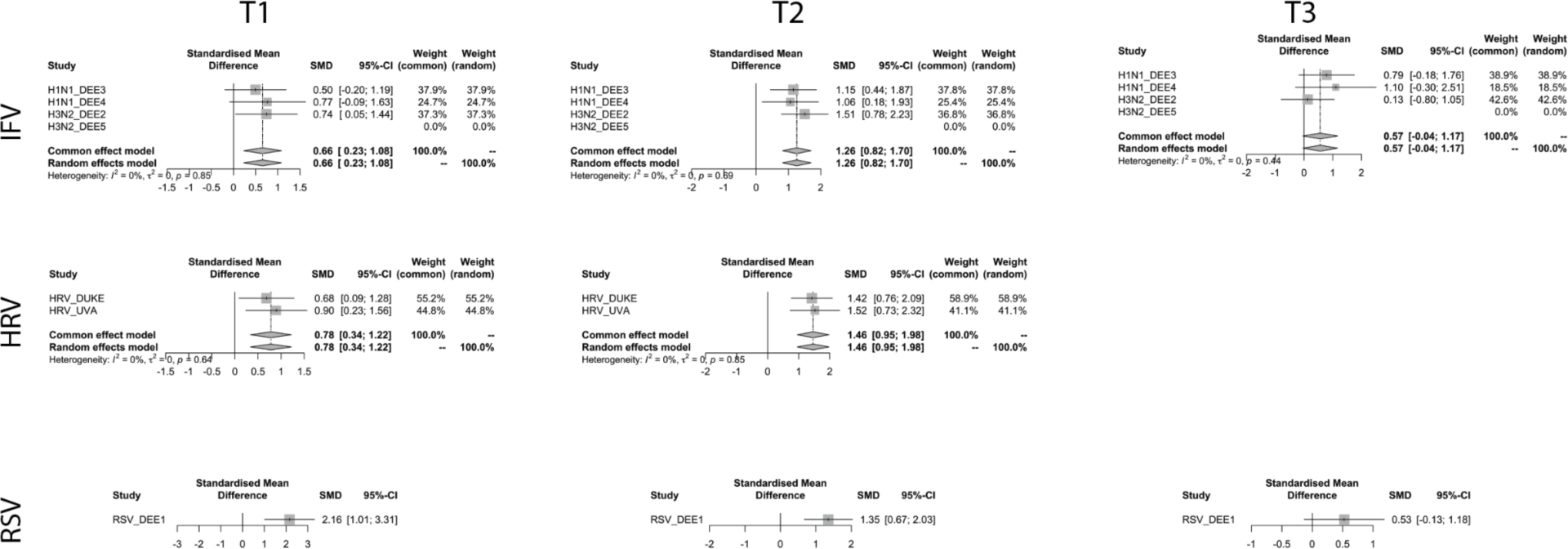
Meta-analysis of Human symptomatic challenge datasets by virus and time category. Meta-analysis of the MVS score across timepoints categories and viral infection in longitudinal human datasets.

**SFig6.**
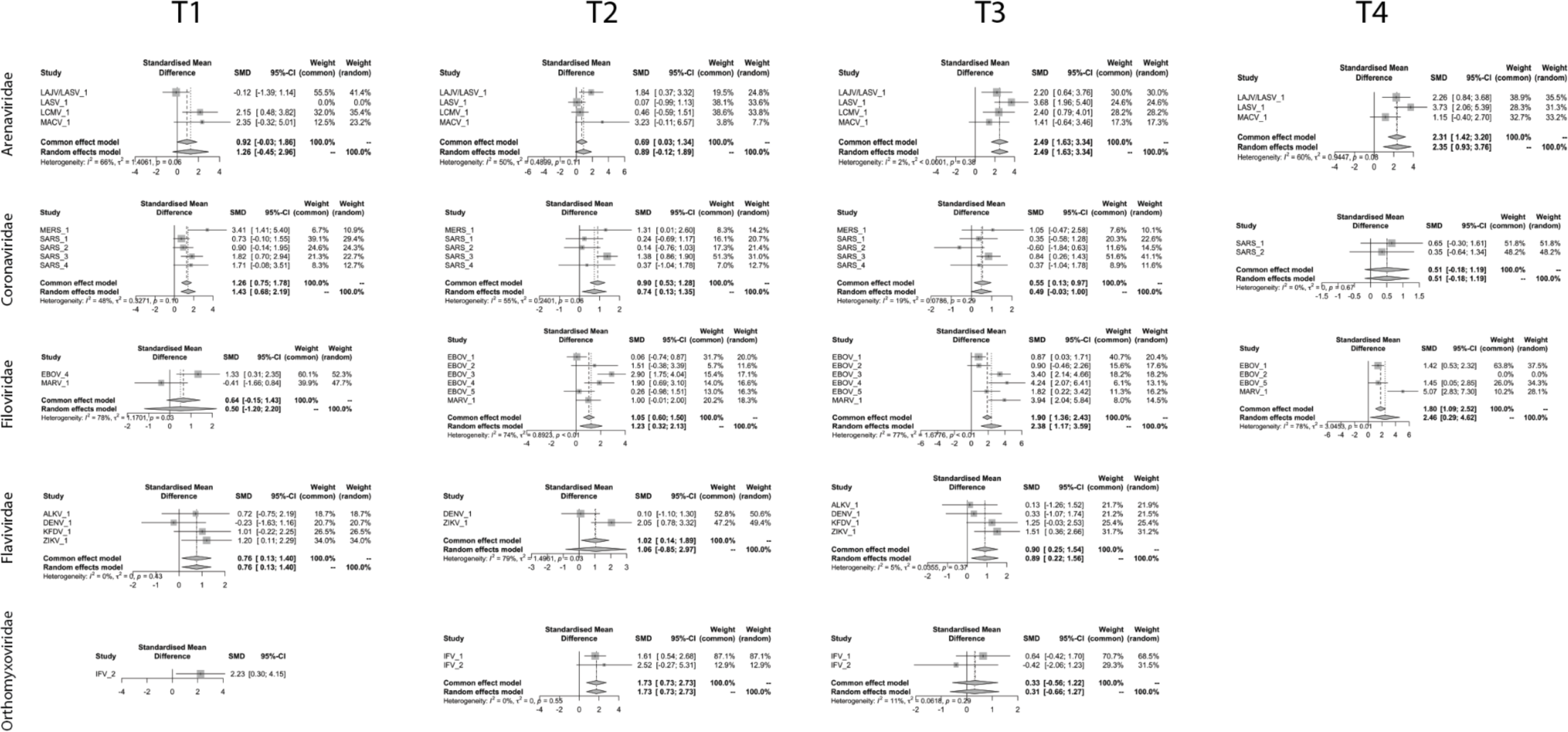
Meta-analysis of NHP datasets by virus and time category. Meta-analysis of the MVS score across timepoints categories and viral infection in macaque datasets.

**SFig7:**
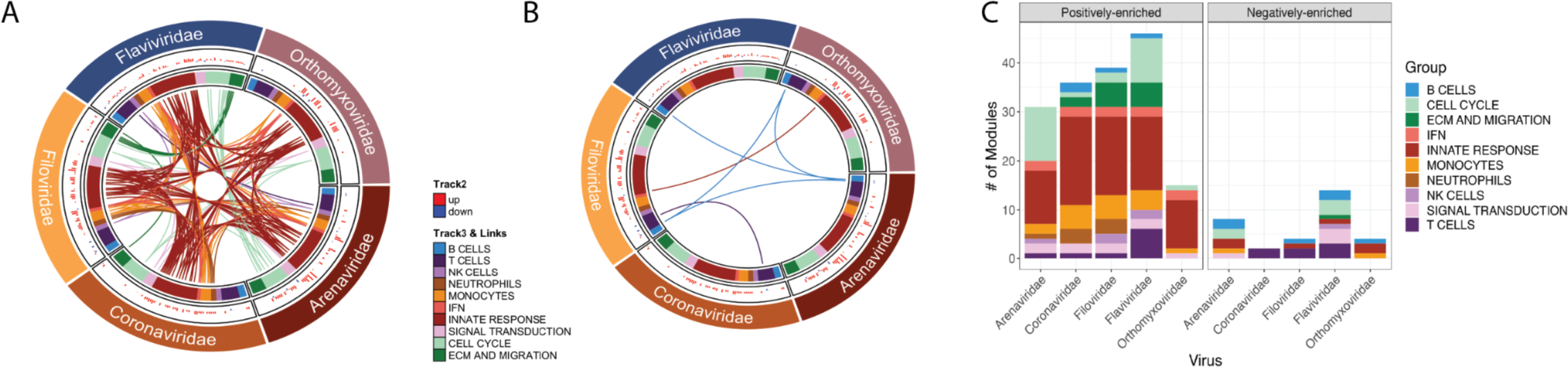
BTM enrichment analysis across DEGs at peak timepoints per virus. (A-B) Circos plots of BTM enrichment analysis across (A) upregulated and (B) downregulated genes at peak infection timepoints in NHP datasets. Each sector represents a viral family, each point in all the tracks represents a BTM that was significant in at least one virus (padj <0.1). Track 2 is a barplot of the geometric mean of the expression of the genes represented by the BTM and plotted where the BTM was significant (padj <0.1). Each color in Track 3 is a granular annotation for each BTM pathway. The inner track connects the same BTM across viral families if they are both (left) positively or (right) negatively enriched. (C) The count of the number of significant modules corresponding to each granular pathway by positive versus negative enichment and by viral family - represented in the barplot.

**SFig8.**
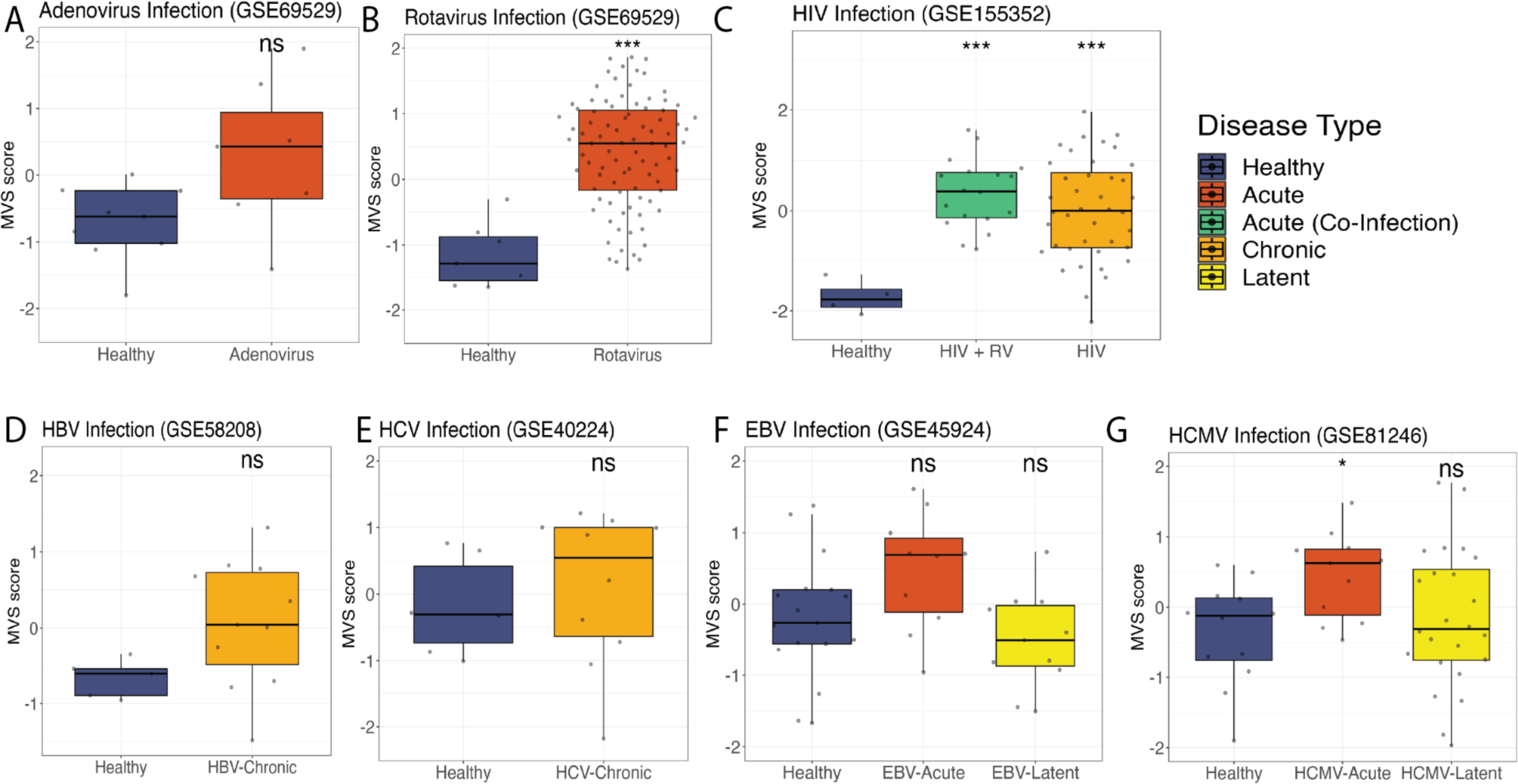
MVS score across human DNA and chronic viruses. (A-G) MVS score in blood samples from healthy control subjects versus patients with (A) Adenovirus infection, (B) Rotavirus infection, (C) acute or latent EBV infection, (D) acute or latent HCMV infection, (E) HIV infection or HIV co-infection with a respiratory virus RV), (F) chronic HBV infection, and (G) chronic HCV infection. (A-G) Significance values were determined using an unpaired Wilcoxon ranked-sum test comparing each condition to healthy samples.. Bonferroni correction for multiple hypothesis testing was applied per subfigure and significance values were assigned by asterisk. Asterisk values across figure are represented as follows: *p value < 0.05, **p value < 0.01, ***p value < 0.001, and ****p value < 0.0001. RV = respiratory virus.

**SFig9:**
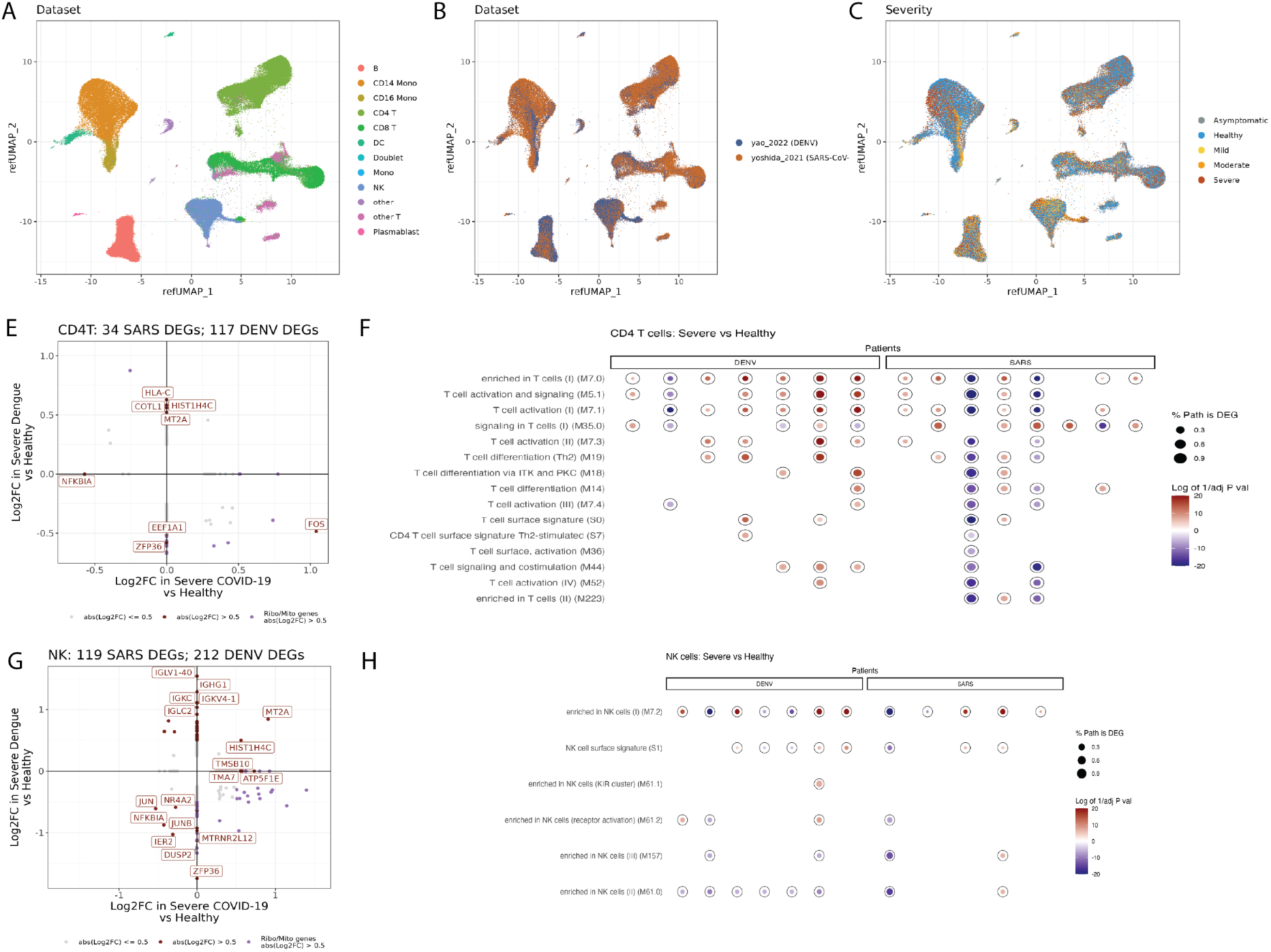
Overview of human scRNA-seq datasets. **(A-C)** UMAP visualization of immune cells from patients colored by **(A)** cell type, **(B)** dataset, and **(C)** patient severity. **(E and G)** Differential gene expression analysis of **(E)** CD4 T cells and **(G)** NK cells across scRNA-seq data from COVID-19 and dengue patients between patients with severe disease compared to healthy controls. **(F and H)** BTM enrichment analysis of differentially expressed genes from **(F)** CD4 T cells and **(H)** NK cells from each severe patient compared to the dataset’s healthy patients.

**Table S3.** Time Series Analysis of MVS Score by Viral Challenge in Humans

| Random effects: | Variance | Std. Deviation | Corr |  |  |
| --- | --- | --- | --- | --- | --- |
| Participant | 0.404 | 0.635 |  |  |  |
| Time | 0.014 | 0.118 | -0.36 |  |  |
| Residual | 0.574 | 0.758 |  |  |  |
| Fixed Effects: | $\beta$ parameter estimate | Std. Error | t value | Pr(>\|t\|) | |
| Intercept | -0.537 | 0.137 | -3.932 | 1.70E-04 | *** |
| Time | 0.904 | 0.059 | 15.443 | < 2e-16 | *** |
| Time x Time | -0.133 | 0.008 | -16.214 | < 2e-16 | *** |
| <u>Virus</u> |  |  |  |  |  |
| Picornaviridae (HRV) | -0.326 | 0.211 | -1.547 | 0.126 |  |
| Pneumoviridae (RSV) | 0.019 | 0.278 | 0.070 | 0.944 |  |
| <u>Virus x Time</u> |  |  |  |  |  |
| Picornaviridae (HRV) x Time | 0.145 | 0.112 | 1.288 | 0.198 |  |
| Pneumoviridae (RSV) x Time | -0.642 | 0.118 | -5.465 | 6.45E-08 | *** |
| <u>Virus x Time x Time</u> |  |  |  |  |  |
| Picornaviridae (HRV) x Time x Time | -0.032 | 0.019 | -1.623 | 0.105 |  |
| Pneumoviridae (RSV) x Time x Time | 0.134 | 0.016 | 8.234 | 5.43E-16 | *** |
| Number of observations: | 1158 |  |  |  |  |
| Number of participants: | 64 |  |  |  |  |
| AIC: | 2918.891 |  |  |  |  |

Mixed effects model using R package lmerTest. Comparison are to Orthomyxoviridae (IFV) challenge. Data from respiratory viral challenge of subjects with symptomatic disease.
Timepoints included were from day 0 to day 7 post-virus challenge.
Signif. codes: 0 '\*\*\*' 0.001 '\*\*' 0.01 '\*' 0.05

## References

1. Rosenberg, R. (2015). Detecting the emergence of novel, zoonotic viruses pathogenic to humans. Cell Mol Life Sci 72, 1115–1125. 10.1007/s00018-014-1785-y.

2. Carrasco-Hernandez, R., Jácome, R., Vidal, Y.L., and León, S.P. de (2017). Are RNA Viruses Candidate Agents for the Next Global Pandemic? A Review. Ilar J 58, 343–358. 10.1093/ilar/ilx026.

3. Morens, D.M., and Fauci, A.S. (2020). Emerging Pandemic Diseases: How We Got to COVID-19. Cell 182, 1077–1092. 10.1016/j.cell.2020.08.021.

4. Kilpatrick, A.M., and Randolph, S.E. (2012). Drivers, dynamics, and control of emerging vector-borne zoonotic diseases. Lancet 380, 1946–1955. 10.1016/s0140-6736(12)61151-9.

5. Domingo, E., and Holland, J.J. (1997). RNA VIRUS MUTATIONS AND FITNESS FOR SURVIVAL. Annu Rev Microbiol 51, 151–178. 10.1146/annurev.micro.51.1.151.

6. Estes, J.D., Wong, S.W., and Brenchley, J.M. (2018). Nonhuman primate models of human viral infections. Nat Rev Immunol 18, 390–404. 10.1038/s41577-018-0005-7.

7. Andres-Terre, M., McGuire, H.M., Pouliot, Y., Bongen, E., Sweeney, T.E., Tato, C.M., and Khatri, P. (2015). Integrated, Multi-cohort Analysis Identifies Conserved Transcriptional Signatures across Multiple Respiratory Viruses. Immunity 43, 1199–1211. 10.1016/j.immuni.2015.11.003.

8. Zheng, H., Rao, A.M., Dermadi, D., Toh, J., Jones, L.M., Donato, M., Liu, Y., Su, Y., Dai, C.L., Kornilov, S.A., et al. (2021). Multi-cohort analysis of host immune response identifies conserved protective and detrimental modules associated with severity across viruses. Immunity 54, 753–768.e5. 10.1016/j.immuni.2021.03.002.

9. Prioritizing diseases for research and development in emergency contexts https://www.who.int/activities/prioritizing-diseases-for-research-and-development-in-emergency-contexts.

10. Reyes, M., Filbin, M.R., Bhattacharyya, R.P., Sonny, A., Mehta, A., Billman, K., Kays, K.R., Pinilla-Vera, M., Benson, M.E., Cosimi, L.A., et al. (2021). Plasma from patients with bacterial sepsis or severe COVID-19 induces suppressive myeloid cell production from hematopoietic progenitors in vitro. Sci Transl Med 13, eabe9599. 10.1126/scitranslmed.abe9599.

11. Reynard, S., Carnec, X., Picard, C., Borges-Cardoso, V., Journeaux, A., Mateo, M., Germain, C., Hortion, J., Albrecht, L., Perthame, E., et al. (2023). A MOPEVAC multivalent vaccine induces sterile protection against New World arenaviruses in non-human primates. Nat Microbiol 8, 64–76. 10.1038/s41564-022-01281-y.

12. Marzi, A., Feldmann, F., Hanley, P.W., Scott, D.P., Günther, S., and Feldmann, H. (2015). Delayed Disease Progression in Cynomolgus Macaques Infected with Ebola Virus Makona Strain. Emerg Infect Dis 21, 1777–1783. 10.3201/eid2110.150259.

13. Maroney, K.J., Pinski, A.N., Marzi, A., and Messaoudi, I. (2021). Transcriptional Analysis of Infection With Early or Late Isolates From the 2013–2016 West Africa Ebola Virus Epidemic Does Not Suggest Attenuated Pathogenicity as a Result of Genetic Variation. Front Microbiol 12, 714817. 10.3389/fmicb.2021.714817.

14. Fukuyama, S., Iwatsuki-Horimoto, K., Kiso, M., Nakajima, N., Gregg, R.W., Katsura, H., Tomita, Y., Maemura, T., Lopes, T.J. da S., Watanabe, T., et al. (2020). Pathogenesis of Influenza A(H7N9) Virus in Aged Nonhuman Primates. J Infect Dis 222, 1155–1164. 10.1093/infdis/jiaa267.

15. Skinner, J.A., Zurawski, S.M., Sugimoto, C., Vinet-Oliphant, H., Vinod, P., Xue, Y., Russell-Lodrigue, K., Albrecht, R.A., García-Sastre, A., Salazar, A.M., et al. (2014). Immunologic characterization of a rhesus macaque H1N1 challenge model for candidate influenza virus vaccine assessment. Clin Vaccine Immunol Cvi 21, 1668–1680. 10.1128/cvi.00547-14.

16. Kotliar, D., Lin, A.E., Logue, J., Hughes, T.K., Khoury, N.M., Raju, S.S., Wadsworth, M.H., Chen, H., Kurtz, J.R., Dighero-Kemp, B., et al. (2020). Single-Cell Profiling of Ebola Virus Disease In Vivo Reveals Viral and Host Dynamics. Cell 183, 1383–1401.e19. 10.1016/j.cell.2020.10.002.

17. Sweeney, T.E., Haynes, W.A., Vallania, F., Ioannidis, J.P., and Khatri, P. (2017). Methods to increase reproducibility in differential gene expression via meta-analysis. Nucleic Acids Res 45, e1–e1. 10.1093/nar/gkw797.

18. Haynes, W.A., Vallania, F., Liu, C., Bongen, E., Tomczak, A., Andres-Terrè, M., Lofgren, S., Tam, A., Deisseroth, C.A., Li, M.D., et al. (2017). EMPOWERING MULTI-COHORT GENE EXPRESSION ANALYSIS TO INCREASE REPRODUCIBILITY. Biocomput 2017 22, 144–153. 10.1142/9789813207813_0015.

19. Ghita, L., Yao, Z., Xie, Y., Duran, V., Cagirici, H.B., Samir, J., Osman, I., Rojas, O.L.A., Sanz, A.M., Sahoo, M.K., et al. (2023). Global and cell type-specific immunological hallmarks of severe dengue progression. Biorxiv, 2022.12.11.519930. 10.1101/2022.12.11.519930.

20. Yoshida, M., Worlock, K.B., Huang, N., Lindeboom, R.G.H., Butler, C.R., Kumasaka, N., Conde, C.D., Mamanova, L., Bolt, L., Richardson, L., et al. (2022). Local and systemic responses to SARS-CoV-2 infection in children and adults. Nature 602, 321–327. 10.1038/s41586-021-04345-x.

21. Lessler, J., Reich, N.G., Brookmeyer, R., Perl, T.M., Nelson, K.E., and Cummings, D.A. (2009). Incubation periods of acute respiratory viral infections: a systematic review. Lancet Infect Dis 9, 291–300. 10.1016/s1473-3099(09)70069-6.

22. Eichner, M., Dowell, S.F., and Firese, N. (2011). Incubation Period of Ebola Hemorrhagic Virus Subtype Zaire. Osong Public Heal Res Perspectives 2, 3–7. 10.1016/j.phrp.2011.04.001.

23. Holmes, G.P., McCormick, J.B., Trock, S.C., Chase, R.A., Lewis, S.M., Mason, C.A., Hall, P.A., Brammer, L.S., Perez- Oronoz, G.I., McDonnell, M.K., et al. (1990). Lassa Fever in the United States. New Engl J Medicine 323, 1120–1123. 10.1056/nejm199010183231607.

24. Wu, Y., Kang, L., Guo, Z., Liu, J., Liu, M., and Liang, W. (2022). Incubation Period of COVID-19 Caused by Unique SARS-CoV-2 Strains. Jama Netw Open 5, e2228008. 10.1001/jamanetworkopen.2022.28008.

25. Jiang, X., Rayner, S., and Luo, M. (2020). Does SARS-CoV-2 has a longer incubation period than SARS and MERS? J. Med. Virol. 92, 476–478. 10.1002/jmv.25708.

26. Virlogeux, V., Yang, J., Fang, V.J., Feng, L., Tsang, T.K., Jiang, H., Wu, P., Zheng, J., Lau, E.H.Y., Qin, Y., et al. (2016). Association between the Severity of Influenza A(H7N9) Virus Infections and Length of the Incubation Period. Plos One 11, e0148506. 10.1371/journal.pone.0148506.

27. Jensen, S., and Thomsen, A.R. (2012). Sensing of RNA Viruses: a Review of Innate Immune Receptors Involved in Recognizing RNA Virus Invasion. J Virol 86, 2900–2910. 10.1128/jvi.05738-11.

28. Yoneyama, M., Kikuchi, M., Natsukawa, T., Shinobu, N., Imaizumi, T., Miyagishi, M., Taira, K., Akira, S., and Fujita, T. (2004). The RNA helicase RIG-I has an essential function in double-stranded RNA-induced innate antiviral responses. Nat Immunol 5, 730–737. 10.1038/ni1087.

29. Andrejeva, J., Childs, K.S., Young, D.F., Carlos, T.S., Stock, N., Goodbourn, S., and Randall, R.E. (2004). The V proteins of paramyxoviruses bind the IFN-inducible RNA helicase, mda-5, and inhibit its activation of the IFN-beta promoter. P Natl Acad Sci Usa 101, 17264–17269. 10.1073/pnas.0407639101.

30. Ramos, H.J., and Gale, M. (2011). RIG-I like receptors and their signaling crosstalk in the regulation of antiviral immunity. Curr Opin Virol 1, 167–176. 10.1016/j.coviro.2011.04.004.

31. Weber, F., Kochs, G., and Haller, O. (2004). Inverse Interference: How Viruses Fight the Interferon System. Viral Immunol 17, 498–515. 10.1089/vim.2004.17.498.

32. Carrat, F., Vergu, E., Ferguson, N.M., Lemaitre, M., Cauchemez, S., Leach, S., and Valleron, A.-J. (2008). Time Lines of Infection and Disease in Human Influenza: A Review of Volunteer Challenge Studies. Am J Epidemiol 167, 775–785. 10.1093/aje/kwm375.

33. Ebola Disease - Diagnosis https://www.cdc.gov/vhf/ebola/diagnosis/index.html#:~:text=Ebola%20virus%20can%20be%20detected,low%20levels%20of%20Ebola%20virus.

34. Salguero, F.J., White, A.D., Slack, G.S., Fotheringham, S.A., Bewley, K.R., Gooch, K.E., Longet, S., Humphries, H.E., Watson, R.J., Hunter, L., et al. (2021). Comparison of rhesus and cynomolgus macaques as an infection model for COVID-19. Nat Commun 12, 1260. 10.1038/s41467-021-21389-9.

35. Coleman, C., Doyle-Meyers, L.A., Russell-Lodrigue, K.E., Golden, N., Threeton, B., Song, K., Pierre, G., Baribault, C., Bohm, R.P., Maness, N.J., et al. (2021). Similarities and Differences in the Acute-Phase Response to SARS-CoV-2 in Rhesus Macaques and African Green Monkeys. Front Immunol 12, 754642. 10.3389/fimmu.2021.754642.

36. Pinski, A.N., Maroney, K.J., Marzi, A., and Messaoudi, I. (2021). Distinct transcriptional responses to fatal Ebola virus infection in cynomolgus and rhesus macaques suggest species-specific immune responses. Emerg Microbes Infec 10, 1320– 1330. 10.1080/22221751.2021.1942229.

37. Cai, C., Tang, Y.-D., Xu, G., and Zheng, C. (2021). The crosstalk between viral RNA- and DNA-sensing mechanisms. Cell Mol Life Sci 78, 7427–7434. 10.1007/s00018-021-04001-7.

38. Fan, Y.M., Zhang, Y.L., Luo, H., and Mohamud, Y. (2022). Crosstalk between RNA viruses and DNA sensors: Role of the cGAS-STING signalling pathway. Rev Med Virol 32, e2343. 10.1002/rmv.2343.

39. Ni, G., Ma, Z., and Damania, B. (2018). cGAS and STING: At the intersection of DNA and RNA virus-sensing networks. Plos Pathog 14, e1007148. 10.1371/journal.ppat.1007148.

40. Neufeldt, C.J., Cerikan, B., Cortese, M., Frankish, J., Lee, J.-Y., Plociennikowska, A., Heigwer, F., Prasad, V., Joecks, S., Burkart, S.S., et al. (2022). SARS-CoV-2 infection induces a pro-inflammatory cytokine response through cGAS-STING and NF-κB. Commun Biology 5, 45. 10.1038/s42003-021-02983-5.

41. Ng, W.C., Kwek, S.S., Sun, B., Yousefi, M., Ong, E.Z., Tan, H.C., Puschnik, A.S., Chan, K.R., Ooi, Y.S., and Ooi, E.E. (2022). A fast-growing dengue virus mutant reveals a dual role of STING in response to infection. Open Biol 12, 220227. 10.1098/rsob.220227.

42. Webb, L.G., and Fernandez-Sesma, A. (2022). RNA viruses and the cGAS-STING pathway: reframing our understanding of innate immune sensing. Curr Opin Virol 53, 101206. 10.1016/j.coviro.2022.101206.

43. Hertzog, J., Zhou, W., Fowler, G., Rigby, R.E., Bridgeman, A., Blest, H.T., Cursi, C., Chauveau, L., Davenne, T., Warner, B.E., et al. (2022). Varicella-Zoster virus ORF9 is an antagonist of the DNA sensor cGAS. Embo J 41, e109217. 10.15252/embj.2021109217.

44. Schmid, S., Mordstein, M., Kochs, G., García-Sastre, A., and tenOever, B.R. (2010). Transcription Factor Redundancy Ensures Induction of the Antiviral State*. J Biol Chem 285, 42013–42022. 10.1074/jbc.m110.165936.

45. Crotta, S., Davidson, S., Mahlakoiv, T., Desmet, C.J., Buckwalter, M.R., Albert, M.L., Staeheli, P., and Wack, A. (2013). Type I and Type III Interferons Drive Redundant Amplification Loops to Induce a Transcriptional Signature in Influenza-Infected Airway Epithelia. Plos Pathog 9, e1003773. 10.1371/journal.ppat.1003773.

46. Fan, C., Wu, Y., Rui, X., Yang, Y., Ling, C., Liu, S., Liu, S., and Wang, Y. (2022). Animal models for COVID-19: advances, gaps and perspectives. Signal Transduct Target Ther 7, 220. 10.1038/s41392-022-01087-8.

47. Kayesh, M.E.H., and Tsukiyama-Kohara, K. (2022). Mammalian animal models for dengue virus infection: a recent overview. Arch Virol 167, 31–44. 10.1007/s00705-021-05298-2.

48. Hanley, K.A., Monath, T.P., Weaver, S.C., Rossi, S.L., Richman, R.L., and Vasilakis, N. (2013). Fever versus fever: The role of host and vector susceptibility and interspecific competition in shaping the current and future distributions of the sylvatic cycles of dengue virus and yellow fever virus. Infect Genetics Evol 19, 292–311. 10.1016/j.meegid.2013.03.008.

49. Goethals, O., Kaptein, S.J.F., Kesteleyn, B., Bonfanti, J.-F., Wesenbeeck, L.V., Bardiot, D., Verschoor, E.J., Verstrepen, B.E., Fagrouch, Z., Putnak, J.R., et al. (2023). Blocking NS3–NS4B interaction inhibits dengue virus in non-human primates. Nature 615, 678–686. 10.1038/s41586-023-05790-6.

50. Chiramel, A.I., Meyerson, N.R., McNally, K.L., Broeckel, R.M., Montoya, V.R., Méndez-Solís, O., Robertson, S.J., Sturdevant, G.L., Lubick, K.J., Nair, V., et al. (2019). TRIM5α Restricts Flavivirus Replication by Targeting the Viral Protease for Proteasomal Degradation. Cell Reports 27, 3269–3283.e6. 10.1016/j.celrep.2019.05.040.

51. Broeckel, R.M., Feldmann, F., McNally, K.L., Chiramel, A.I., Sturdevant, G.L., Leung, J.M., Hanley, P.W., Lovaglio, J., Rosenke, R., Scott, D.P., et al. (2021). A pigtailed macaque model of Kyasanur Forest disease virus and Alkhurma hemorrhagic disease virus pathogenesis. Plos Pathog 17, e1009678. 10.1371/journal.ppat.1009678.

52. Fathi, N., and Rezaei, N. (2020). Lymphopenia in COVID-19: Therapeutic opportunities. Cell. Biol. Int. 44, 1792–1797. 10.1002/cbin.11403.

53. Diao, B., Wang, C., Tan, Y., Chen, X., Liu, Y., Ning, L., Chen, L., Li, M., Liu, Y., Wang, G., et al. (2020). Reduction and Functional Exhaustion of T Cells in Patients With Coronavirus Disease 2019 (COVID-19). Front Immunol 11, 827. 10.3389/fimmu.2020.00827.

54. Kahan, S.M., Wherry, E.J., and Zajac, A.J. (2015). T cell exhaustion during persistent viral infections. Virology 479, 180–193. 10.1016/j.virol.2014.12.033.

55. Iampietro, M., Younan, P., Nishida, A., Dutta, M., Lubaki, N.M., Santos, R.I., Koup, R.A., Katze, M.G., and Bukreyev, A. (2017). Ebola virus glycoprotein directly triggers T lymphocyte death despite of the lack of infection. Plos Pathog 13, e1006397. 10.1371/journal.ppat.1006397.

56. John, A.L.St., and Rathore, A.P.S. (2019). Adaptive immune responses to primary and secondary dengue virus infections. Nat Rev Immunol 19, 218–230. 10.1038/s41577-019-0123-x.

57. Tian, Y., Seumois, G., De-Oliveira-Pinto, L.M., Mateus, J., Mata, S.H. la, Kim, C., Hinz, D., Goonawardhana, N.D.S., Silva, A.D. de, Premawansa, S., et al. (2019). Molecular Signatures of Dengue Virus-Specific IL-10/IFN-γ Co-producing CD4 T Cells and Their Association with Dengue Disease. Cell Reports 29, 4482–4495.e4. 10.1016/j.celrep.2019.11.098.

58. Grifoni, A., Voic, H., Yu, E.D., Mateus, J., Fung, K.M.Y., Wang, A., Seumois, G., Silva, A.D.D., Tennekon, R., Premawansa, S., et al. (2022). Transcriptomics of Acute DENV-Specific CD8+ T Cells Does Not Support Qualitative Differences as Drivers of Disease Severity. Nato Adv Sci Inst Se 10, 612. 10.3390/vaccines10040612.

59. Tian, Y., Grifoni, A., Sette, A., and Weiskopf, D. (2019). Human T Cell Response to Dengue Virus Infection. Front Immunol 10, 2125. 10.3389/fimmu.2019.02125.

60. Hatch, S., Endy, T.P., Thomas, S., Mathew, A., Potts, J., Pazoles, P., Libraty, D.H., Gibbons, R., and Rothman, A.L. (2011). Intracellular Cytokine Production by Dengue Virus–specific T cells Correlates with Subclinical Secondary Infection. J Infect Dis 203, 1282–1291. 10.1093/infdis/jir012.

61. Screaton, G., Mongkolsapaya, J., Yacoub, S., and Roberts, C. (2015). New insights into the immunopathology and control of dengue virus infection. Nat Rev Immunol 15, 745–759. 10.1038/nri3916.

62. Patro, R., Duggal, G., Love, M.I., Irizarry, R.A., and Kingsford, C. (2017). Salmon provides fast and bias-aware quantification of transcript expression. Nat Methods 14, 417–419. 10.1038/nmeth.4197.

63. Soneson, C., Love, M.I., and Robinson, M.D. (2016). Differential analyses for RNA-seq: transcript-level estimates improve gene-level inferences. F1000Research 4, 1521. 10.12688/f1000research.7563.2.

64. Love, M.I., Huber, W., and Anders, S. (2014). Moderated estimation of fold change and dispersion for RNA-seq data with DESeq2. Genome Biol 15, 550. 10.1186/s13059-014-0550-8.

65. Buturovic, L., Zheng, H., Tang, B., Lai, K., Kuan, W.S., Gillett, M., Santram, R., Shojaei, M., Almansa, R., Nieto, J.Á., et al. (2022). A 6-mRNA host response classifier in whole blood predicts outcomes in COVID-19 and other acute viral infections. Sci Rep-uk 12, 889. 10.1038/s41598-021-04509-9.

66. Liu, Y.E., Saul, S., Rao, A.M., Robinson, M.L., Rojas, O.L.A., Sanz, A.M., Verghese, M., Solis, D., Sibai, M., Huang, C.H., et al. (2022). An 8-gene machine learning model improves clinical prediction of severe dengue progression. Genome Med 14, 33. 10.1186/s13073-022-01034-w.

67. Simmons, C.P., Popper, S., Dolocek, C., Chau, T.N.B., Griffiths, M., Dung, N.T.P., Long, T.H., Hoang, D.M., Chau, N.V., Thao, L.T.T., et al. (2007). Patterns of Host Genome—Wide Gene Transcript Abundance in the Peripheral Blood of Patients with Acute Dengue Hemorrhagic Fever. J. Infect. Dis. 195, 1097–1107. 10.1086/512162.

68. Kwissa, M., Nakaya, H.I., Onlamoon, N., Wrammert, J., Villinger, F., Perng, G.C., Yoksan, S., Pattanapanyasat, K., Chokephaibulkit, K., Ahmed, R., et al. (2014). Dengue Virus Infection Induces Expansion of a CD14+CD16+ Monocyte Population that Stimulates Plasmablast Differentiation. Cell Host Microbe 16, 115–127. 10.1016/j.chom.2014.06.001.

69. Popper, S.J., Gordon, A., Liu, M., Balmaseda, A., Harris, E., and Relman, D.A. (2012). Temporal Dynamics of the Transcriptional Response to Dengue Virus Infection in Nicaraguan Children. PLoS Neglected Trop. Dis. 6, e1966. 10.1371/journal.pntd.0001966.

70. Long, H.T., Hibberd, M.L., Hien, T.T., Dung, N.M., Ngoc, T.V., Farrar, J., Wills, B., and Simmons, C. (2009). Patterns of gene transcript abundance in the blood of children with severe or uncomplicated dengue highlight differences in disease evolution and host response to dengue virus infection. J. Infect. Dis. 199, 537–546. 10.1086/596507.

71. Sweeney, T.E., Wong, H.R., and Khatri, P. (2016). Robust classification of bacterial and viral infections via integrated host gene expression diagnostics. Sci. Transl. Med. 8, 346ra91. 10.1126/scitranslmed.aaf7165.

72. Li, S., Rouphael, N., Duraisingham, S., Romero-Steiner, S., Presnell, S., Davis, C., Schmidt, D.S., Johnson, S.E., Milton, A., Rajam, G., et al. (2014). Molecular signatures of antibody responses derived from a systems biological study of 5 human vaccines. Nat Immunol 15, 195–204. 10.1038/ni.2789.

73. Wilk, A.J., Lee, M.J., Wei, B., Parks, B., Pi, R., Martínez-Colón, G.J., Ranganath, T., Zhao, N.Q., Taylor, S., Becker, W., et al. (2021). Multi-omic profiling reveals widespread dysregulation of innate immunity and hematopoiesis in COVID-19. J Exp Med 218, e20210582. 10.1084/jem.20210582.

74. Hagan, T., Gerritsen, B., Tomalin, L.E., Fourati, S., Mulè, M.P., Chawla, D.G., Rychkov, D., Henrich, E., Miller, H.E.R., Diray-Arce, J., et al. (2022). Transcriptional atlas of the human immune response to 13 vaccines reveals a common predictor of vaccine-induced antibody responses. Nat Immunol 23, 1788–1798. 10.1038/s41590-022-01328-6.

75. Hao, Y., Hao, S., Andersen-Nissen, E., Mauck, W.M., Zheng, S., Butler, A., Lee, M.J., Wilk, A.J., Darby, C., Zager, M., et al. (2021). Integrated analysis of multimodal single-cell data. Cell 184, 3573–3587.e29. 10.1016/j.cell.2021.04.048.

